# Electron bifurcation arises from emergent features of multicofactor enzymes

**DOI:** 10.1101/2025.04.30.651465

**Authors:** Anna Wójcik-Augustyn, Łukasz Bujnowicz, Artur Osyczka, Marcin Sarewicz

## Abstract

Quinone-based electron bifurcation (EB) catalyzed by cytochrome *bc*_1_ (cyt*bc*_1_) plays a critical role in maximizing efficiency of biological energy conversion. The canonical EB model (CEB), grounded in equilibrium redox potentials, dictates the order of EB steps with initial endergonic reduction of high-potential iron-sulfur cluster (2Fe2S) by quinol followed by exergonic reduction of low-potential heme *b*_L_ (*b*_L_) by semiquinone (SQ). However, this concept falls short in explaining several experimental observations, including intermediate semiquinone spin-coupled to 2Fe2S (SQ-2Fe2S^red^) and the absence of short-circuiting. Presented here DFT calculations on large cluster models of cyt*bc*_1_, encompassing both 2Fe2S and *b*_L_, identified location of donor (HOMO) and acceptor (LUMO) orbitals along with the previously not considered microstates to reveal that EB is an emergent property of an integrated system of redox cofactors where transient charge separations dynamically modulate electron affinities. In this system, electron transfer initiates preferentially toward *b*_L_, indicating a departure from the conventional sequence proposed by CEB. Based on this finding, we introduce an EMBER (EMergent B_L_-first Electron Routing) model of EB and demonstrate that its assumptions are supported by electron paramagnetic resonance spectroscopy data. Unlike CEB, EMBER proposes a relatively flat energy profile for EB that accommodates stable SQ-2Fe2S^red^ and explains suppression of short-circuits without additional assumptions. It highlights the importance of state-dependent electrostatic interactions in shaping electron transfer pathways. In general, the concept of emergence inherent to EMBER offers a mechanistic framework applicable to a broad range of multi-cofactor redox enzymes beyond cyt*bc*_1_.

## Introduction

The quinone-based electronic bifurcation (EB) is a crucial process that maximizes the efficiency of energy conversion during respiration and photosynthesis^1–5^. Although flavin-based EB has also been identified in some enzymes, the present work focuses on the mechanism of EB catalyzed by cytochromes (cyt) *bc* of the Rieske/*b* family, including cytochrome *bc*_1_ (cyt*bc*_1_) in bacteria, mitochondrial complex III, and cyt*b*_6_*f* in plants^2,6,7^. These enzymes share a similar homodimeric structure across all organisms, exhibiting nearly identical core compositions (Fig. 1A) and spatial arrangements of redox cofactors (Fig. 1B) as well as catalytic sites for quinol (QH_2_) oxidation (Q_o_) and quinone (Q) reduction (Q_i_)^2,8,9^. In each monomer, the cofactors are branched into two chains: low- and high-potential, with Q_o_ located between them (Fig. 1C). Upon binding to this site, QH_2_ becomes flanked by two redox cofactors: the Rieske iron–sulfur cluster (2Fe2S) from the high-potential chain and heme *b*_L_ (*b*_L_) from the low-potential chain.

**Figure 1.**
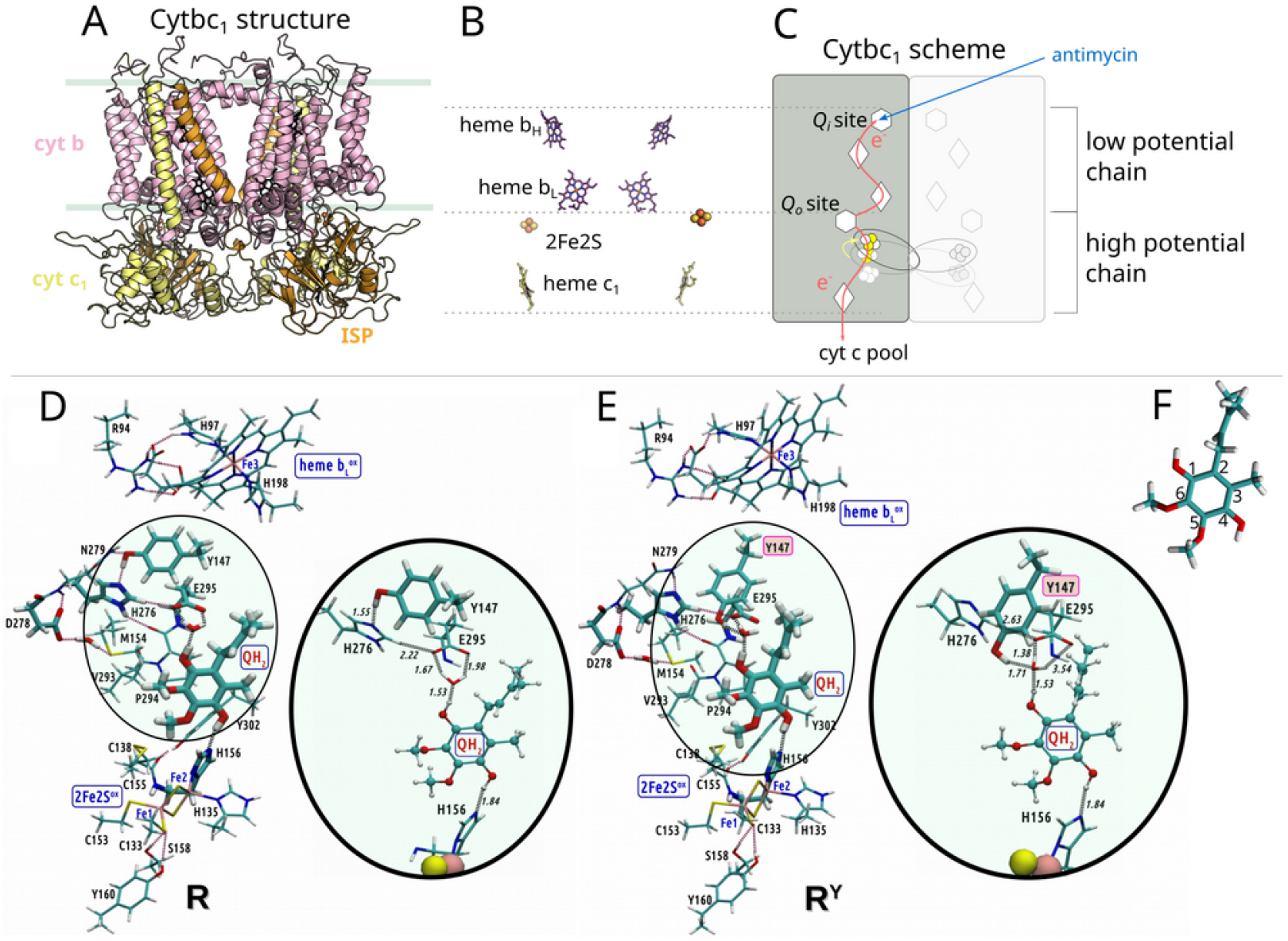
Overview of the structural elements of cyt*bc*_1_. A) The crystallographic model of the cyt*bc*_1_ dimer from *Rhodobacter capsulatus* (PDB ID: 1ZRT). Each monomer consists of 3 catalytic subunits: cytochrome *b* (cyt*b*), cytochrome (cyt*c*_1_) and iron-sulfur protein (ISP). B) The spatial arrangement of redox-active cofactors: hemes *b*_L_, *b*_H_ of cyt*b*, 2FeS of ISP, heme *c*_1_ of cyt*c*_1_. C) General pathways for electrons (*red arrows*) released upon oxidation of QH_2_ at Q_o_ (*white hexagon*). EB directs one electron to high potential chain (consisting of 2Fe2S, *c*_1_ and *c*) and one electron to the low potential chain (consisting of *b*_L_, *b*_H_ and Q_i_). *White diamonds* denote hemes, *circles* denote 2Fe2S. Yellow arrow indicates movement of the head domain of ISP (ISP-HD) containing 2Fe2S between cyt*b* and cyt*c*_1_. The blue arrow shows where inhibitor antimycin binds. For simplicity, the scheme shows eT in one monomer. D) and E) show optimized initial structures of QM model with two possible conformations of cyt *b*:Y147: (D) **R** with Y147 directed toward cyt*b*:H276, and (E) **R**^**Y**^ with Y147 directed toward the bound QH_2_. Fragments of the models, highlighting the different conformations of the Y147 side chain, are magnified. F) The quinone moiety with the carbon atom numbering of the ring indicated.

In cyt*bc*_1_, EB relies on the two-electron oxidation of QH_2_, with each electron diverging into a different direction by transfer to distinct electron acceptors: oxidized *b*_L_ (*b*_L_^ox^) and oxidized 2Fe2S (2Fe2S^ox^):

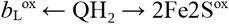

A one-to-one correlation between the number of electrons transferred from QH_2_ to *b* _L_^ox^ and to 2Fe2S^ox^ is expected to ensure maximal energy conservation efficiency, while preventing direct or quinone-mediated electron exchange between *b*_L_ and 2Fe2S. Although this process appears conceptually simple, it has never been reproduced in synthetic systems^10^.

The general framework for EB mechanism, originally proposed by Mitchell, was based on the formalism of equilibrium electrochemistry, i.e. on the equilibrium redox midpoint potentials ^11,12^. It describes EB as two consecutive one-electron steps, with the Q/QH_2_ redox reaction divided into two half-reactions involving the SQ/QH_2_ and Q/SQ redox couples (where SQ and Q denote semiquinone and quinone, respectively). These couples are proposed to differ significantly in their redox midpoint potentials (*E*_m_), with the SQ/QH_2_ couple being very high-potential (>400 mV) and the Q/SQ couple being very low-potential (∼−200 mV)^13,14^. This suggests that EB is driven by large (more than 600 mV) differences between these couples^14^. Accordingly, EB is assumed to begin with an uphill oxidation of QH_2_ to SQ by 2Fe2S (*E*_m_^2Fe2S^ ∼300 mV), followed by downhill oxidation of SQ to Q by *b*_L_ (*E*_m_^*b*L^ ∼−120 mV) and the first uphill step is generally considered as a rate limiting step in the cyt*bc*_1_ catalysis. This specific sequence of QH_2_ oxidation, first by 2Fe2S^ox^ and then by *b*_L_^ox^, derived from consideration of *E*_m_ values for the immediate acceptors (i.e. 2Fe2S and *b*_L_) and the Q/SQ/QH_2_ triad, has become the canonical EB mechanism and the basis for interpreting both experimental and computational studies. Hereafter, this model is referred to as the canonical EB mechanism (CEB).

Several important consequences follow from CEB. First, SQ is expected to be very unstable and transient, as the first step (oxidation of QH_2_ to SQ by 2Fe2S) is endoergic, while the second (oxidation of SQ to Q by *b*_L_) is exoergic^14^. Consequently, SQ is presumed to be highly unstable and potentially a substrate for superoxide radical formation due to low *E*_m_ of Q/SQ couple^15^. This inherent instability may explain the long-standing challenges associated with the direct detection of SQ at the Q_o_ site. Second, considering the *E*_m_ values of *b*_L_, 2Fe2S, and the Q/SQ/QH_2_ triad, the overall reaction should, under certain conditions, be susceptible to internal, quinone-mediated electron exchange between the chains, that is, quinone-mediated electron transfer from *b*_L_ ^red^ to 2Fe2S^ox 2,4^. This issue becomes especially problematic in understanding the inhibition of cyt*bc*_1_ by antimycin^16,17^. According to CEB, inhibition by antimycin should result in the reduction of both *b*_H_ and *b*_L_. Subsequent oxidation of QH_2_ by 2Fe2S^ox^ would then be followed by thermodynamically favorable reduction of SQ back to QH_2_ by eT from *b* _L_ ^red^. As a consequence, EB would be bypassed, with both electrons from QH_2_ flowing exclusively through the high-potential chain. This possible reaction is referred to as short circuit reaction^4,14^. However, so far such events have remained experimentally undetectable. On the other hand, under non-equilibrium conditions in the presence of antimycin, relatively large amounts of SQ spin-coupled to 2Fe2S^red^ (SQ–2Fe2S^red^) and small amounts of SQ free radical signal have been detected^18–22^. Yet, the roles of these species in EB remain poorly understood. While the presence of the SQ signal can still be explained within the framework of CEB, the problem arises when considering the substantial population of SQ spin-coupled to reduced 2Fe2S (SQ–2Fe2S^red^). These unresolved observations and lack of expected short circuits have long hindered full mechanistic understanding of EB in cyt*bc*_1_.

The issue associated with the inhibitory effect of antimycin is particularly important, considering the fact that this inhibitor has been widely used for years in experiments performed to understand the mechanism of EB. Simple models based solely on *E*_m_s face problems in explaining the inhibition of QH_2_ oxidation and the lack of internal short circuits. Over the years, various models have been proposed to overcome these problems, yet the aforementioned issues have never been satisfactorily addressed^2^. Consequently, several, often mutually exclusive, so-called gating mechanisms of EB coexist^14,23–29^, raising the concern that a fundamental principle underlying efficient QH_2_ oxidation by cyt*bc*_1_ might still be missing and that a complete description of Q_o_ catalysis has yet to be formulated^5^.

Here, we employed quantum mechanical (QM) calculations to analyze EB in cyt*bc*_1_ at the level of the electronic function, allowing us to follow the progress of the key steps of the reaction. We also aimed to explain the mechanism of formation and the role of SQ–2Fe2S^red^ in suppressing short circuits and the mechanism of antimycin inhibition of QH_2_ oxidation at the Q_o_ site of cyt*bc*_1_ from purple bacteria *Rhodobacter capsulatus*. To this end, we performed density functional theory (DFT) analyses on models containing both *b*_L_ and 2Fe2S cofactors as inseparable components of the EB reaction. To the best of our knowledge, this approach has not been previously applied in studies of EB mechanism in cyt*bc*_1_ and was chosen to avoid or at least minimize the artificial selection of reaction sequence inherent in models that isolate individual one-electron reactions. Our *ab initio* approach lead us to demonstration that all redox-active elements act as an integrated system of cofactors, whose behavior cannot be reproduced by two separate models comprising the quinone molecule and either *b*_L_ or 2Fe2S. In particular, we show that the initiation of EB deviates significantly from CEB and proceeds through previously unconsidered microstates. Based on the calculations, we developed a model of EB and analyzed its predictions with existing data and the results of newly performed experiments. We believe that these results help reconcile the apparent discrepancies between reports detecting semiquinone intermediates within the Q_o_ site of cyt*bc*_1_ and cyt*b*_6_*f*, while also explaining the absence of short circuiting and the mechanism of antimycin inhibition. If our model correctly describes the EB mechanism, it also underscores the potential limitations of using equilibrium-determined *E*_m_ values to construct each step of the reaction. Rather charge separations, resulting from eT and pT within the protein interior, can modify interactions (Coulombic, for example) between cofactors, leading to state-dependent changes in their electron affinities.

Although our proposition represents a new hypothesis that requires extensive future validation, it offers a unified explanation for several long-standing challenges. We propose that this model provides mechanistic insight into the efficiency of EB and may guide the design of synthetic systems.

## Materials and Methods

### QM model of the Q_o_ active site

The cluster model of the Q_o_ active site was constructed based on the crystallographic structure of cyt*bc*_1_ from *R. capsulatus* (PDB code: 1ZRT)^30^. Stigmatellin bound at the Q_o_ site was replaced with ubiquinol, preserving the position of the isoprene chain, and two water molecules were added based on molecular dynamics simulations^31^. The structure containing stigmatellin was selected as the model for constructing the active state **R**, following suggestions that stigmatellin at Q_o_ mimics the enzyme–substrate complex between quinol and 2Fe2S^ox 32^ consisted of 399 atoms and included the following elements: *b*_L_, ubiquinol (with its hydrophobic tail truncated to one isoprene unit), two water molecules, the 2Fe2S cluster, eight residues from chain E (C133, H135, C138, C153, C155, H156, S158, Y160), and twelve residues from chain P (R94, H97, Y147, M154, H198, H276, D278, N279, V293, P294, E295, Y302). The residues from chain E (C133, H135, C138, C153, S158, Y160) and chain P (R94, H97, Y147, M154, H198, H276, Y302) were included with their amino and carbonyl moieties of the protein backbone replaced by hydrogen atoms. The protein fragments C155–H156 (chain E) and D278–N279, V293–P294–E295 (chain P) were included along with the peptide bonds connecting them, with the corresponding CO and NH groups of the protein backbone substituted by hydrogen atoms (i.e., for C155, D278, and V293 the NH groups were replaced, and for H156, N279, and E295 the CO groups were replaced).

Protonation states of the residues were assigned using program Propka 3.1 ^33,34^, validated by PypKa 2.10.0 ^35^. For the H156 residue, which coordinates the 2Fe2S cluster and is hypothesized to act as a direct proton acceptor during QH_2_ oxidation, two protonation states were tested. In the first state, the H156 ligand was deprotonated and ready to accept a proton, while in the second state, H156 was protonated at the N_τ_ atom. To maintain the rigidity of the protein backbone, geometry optimizations of all structures considered in the present study were performed with constraints imposed on the hydrogen atoms replacing protein backbone moieties and on the carbon atoms that are part of the protein backbone.

### QM model of the active Q_i_ active site

The QM models of the Q_i_ site were constructed based on the crystallographic structure of cyt*bc*_1_ from *Bos taurus* with ubiquinone bound at the Q_i_ site (PDB code: 1NTZ)^36^. This structure was chosen for several reasons. It includes the substrate bound at the Q_i_ site and its quality is sufficient for geometry optimization of large models using QM methods. Also it contains resolved positions of water molecules which, according to the literature, directly participate in the ubiquinone reduction reaction; and its Q_i_ site shows high structural similarity^37^. When comparing the Q_i_ sites from *B. taurus* and *R. capsulatus*, subtle differences can be observed, but they do not involve the residues directly participating in the reaction. The cluster model of the Q_i_ site consisted of 334 atoms and included the following components: heme *b*_H_, ubiquinone with its hydrophobic tail truncated to one isoprene unit (as in the model of the Q_o_ site), four water molecules (in the same positions as in the 1NTZ structure), and thirteen protein residues: I27^*Bt*^/L41^*Rc*^ (*Bt* and *Rc* in superscript denote *Bos taurus* or *R. capsulatus*, respectively), W31^*Bt*^/W45^*Rc*^, H97^*Bt*^/H111^*Rc*^, R100^*Bt*^/R114^*Rc*^, H196^*Bt*^/H212^*Rc*^), L200^*Bt*^/F216^*Rc*^), H201^*Bt*^/H217^*Rc*^, S205^Bt^/N221^*Rc*^, N206^*Bt*^/N222^*Rc*^, F220^*Bt*^/F244^*Rc*^, Y224^*Bt*^/F248^*Rc*^, K227^*Bt*^/K251^*Rc*^, D228^*Bt*^/D252^*Rc*^. In the QM model of the Q_i_ site, the amino and carbonyl groups of the protein backbone were replaced by hydrogen atoms in the following residues: I27, W31, H97, R100, H196, L200, H201, and Y224. In F220, the Cβ carbon was replaced by a hydrogen atom. The fragments S205–N206 and K227–D228 were included in the model with their peptide bonds preserved. Accordingly, only the NH backbone moieties of S205 and K227, and the CO backbone moieties of N206 and D228 were substituted with hydrogen atoms.

Protonation states of the residues were assigned using program Propka 3.1^33,34^ and validated by PypKa 2.10.0^35^. The geometry optimization of intermediates involved in the ubiquinone reduction was perfomed with the constraints imposed on atoms derived from protein backbone, including hydrogen atoms replacing any moieties present in the whole protein structure.

### QM methods

Due to the very large cluster models (∼400 atoms) used in this study, computational limitations rendered vibrational analysis and full transition-state characterization unfeasible^38,39^, therefore, constrained TS-like structures (pre-TS^*^) were employed to provide upper-bound estimates of the transition-state energies. All QM computations were performed using the Gaussian 16 program^40^. The geometries of all stationary points were optimized using DFT with the B3LYP hybrid exchange–correlation functional and Grimme’s D3 dispersion correction with Becke– Johnson damping^41,42^. Geometry optimizations were performed with the double-ζ basis set def2-SVP, while the final energies of the intermediates were computed with the triple-ζ basis set def2-TZVP, within a polarizable continuum model (PCM) characterized by a dielectric constant of 4.0 and a probe radius of 1.4 Å, to mimic the electrostatic effects of the protein environment^43^. The dielectric constant was set to 4 to reflect the highly hydrophobic environment of the Q_o_ sites and the surroundings in cyt*bc*_1_ ^44,45^. These methods have been previously validated for computational studies of active sites involving iron–sulfur clusters and hemes^46,47^.

Since the **P3** structure relaxes to **I2** during geometry optimization with doublet multiplicity in our model, in which *b*_H_ is not explicitly included, its geometry was optimized in the quartet multiplicity state, and the final energy was computed for the doublet state. The presence of the reduced *b*_H_ (*b*_H_^red^) is expected to decrease electron affinity of *b*_L_ due to electrostatic interactions and should stabilize **P3** state in respect to **I2**. To overcome limitations of the optimization algorithm associated with geometry optimization of shallow minimum state of **P3** (in dublet multiplicity), the quartet state was used to optimize geometry of **P3**. The results revealed that energy obtained for quartet and doublet for geometry optimized for quartet multiplicity are identical. Thus, for the spin orientations of *b*_L_^ox^ considered (α, β), the energy of the **P3** state, which corresponds to ferromagnetic coupling between SQ and the reduced 2Fe2S cluster, remains essentially unchanged. This is consistent with the observation that at temperature > 20 K, CW EPR spectra of SQ-2Fe2S^red^ still remains unaffected despite a very fast spin-lattice relaxation rate (>> 4×10^6 s-1^) of iron ion in heme *b*_L_^18,48^.

### Material used for experimental sample preparation

Equine cytochrome *c* (cyt*c*), 2,3-dimethoxy-5-decyl-6-methyl-1,2-benzoquinone (DB), and antimycin were purchased from Sigma-Aldrich (St. Louis, MO, USA). Cyt*bc*_1_ was isolated from *Rhodobacter capsulatus* grown under semiaerobic conditions, as described previously ^49^. The isolated cyt*bc*_1_ solution was dialyzed against reaction buffer composed of 50 mM bicine (pH 8.0), 100 mM NaCl, and 0.01% (w/w) dodecylmaltoside (DDM). DB was reduced to its hydroquinone form (DBH_2_) using sodium borohydride. Iron(III) citrate, used as an internal standard for sample packing efficiency, was prepared immediately before the experiment by adding FeCl_3_ to a tenfold molar excess of citric acid.

### Freeze-quench experiments

Non-equilibrium samples of cyt*bc*_1_ were prepared as described previously^50^. Briefly, freeze-quenched samples were generated using a Biologic SFM-300 stopped-flow mixer equipped with an MPS-70 programmable syringe controller and electron paramagnetic resonance (EPR) freeze– quench accessories. One syringe contained a mixture of cyt*bc*_1_ (50 µM), antimycin (250 µM), cyt*c* (370 µM), and Fe^3+^ citrate (10 µM). The second syringe contained a solution of DBH _2_ (390 µM) in bicine buffer. The reaction was initiated by mixing the contents of the two syringes in a 1:1 volume ratio. The mixture was incubated in a delay line for a programmed period and then injected into an isopentane bath cooled to 170 K. The frozen droplets were collected in an EPR tube and measured immediately after preparation. Different occupancies of the Q_o_ site by quinone radical states were achieved by modulating the sample incubation time.

### Double mixing experiments

For the double mixing experiments shown, we used our home-built multichannel pulse high-current drivers and a custom mixer with a rotary valve, as described previously^51,52^. Cyt*bc*_1_ was prepared at a concentration of 80 µM in bicine buffer (as described above) and supplemented with 100 µM antimycin. Prior to mixing, portions of potassium ferricyanide were added, and the oxidation level of cyt*c*_1_ was continuously monitored spectrophotometrically. Addition of the oxidant was continued until 90% of heme *c*_*1*_ was oxidized. Synthetic ubiquinol (DBH_2_) and equine cyt*c* were prepared in bicine buffer at concentrations of 180 µM and 300 µM, respectively. Syringes 1 and 2 contained 125 µL of cyt*bc*_1_ solution and 125 µL of DBH_2_ solution, respectively. The solutions were mixed and incubated for 500 ms in syringe 3 to allow the enzyme to reach state **P3**. Then, 250 µL of the cyt*bc*_1_ + DBH_2_ mixture was combined with 250 µL of cyt*c* solution and aged for an additional 200 ms. Finally, the mixture containing cyt*bc*_1_, DBH_2_, and cyt*c* was injected into cold isopentane to stop the reaction. This procedure was repeated twice using independently isolated cyt*bc*_1_ preparations, and each experiment was performed in triplicate.

**Table 1.**
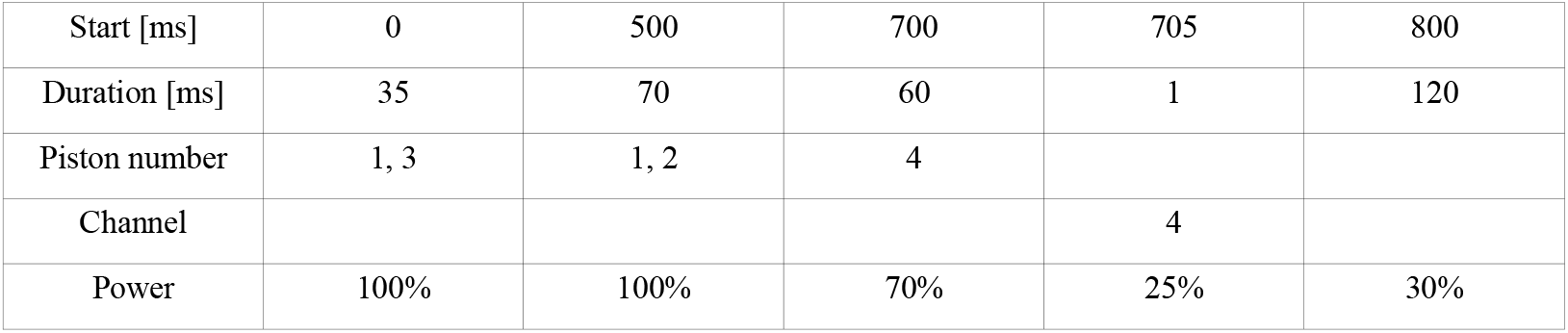
The pattern of pulses used for the multichannel pulse high-current drivers.

|  |  |  |  |  |  |
| --- | --- | --- | --- | --- | --- |
| Start [ms] | 0 | 500 | 700 | 705 | 800 |
| Duration [ms] | 35 | 70 | 60 | 1 | 120 |
| Piston number | 1, 3 | 1, 2 | 4 |  |  |
| Channel |  |  |  | 4 |  |
| Power | 100% | 100% | 70% | 25% | 30% |

### EPR measurements

All EPR measurements were performed using a Bruker Elexsys E580 spectrometer (Bruker, Billerica, MA, USA). X-band continuous wave EPR spectra were measured using a SHQE0511 resonator and an ESR900 cryostat (Oxford Instruments, Abingdon, UK). The helium temperature was maintained using a closed-cycle Stinger cryocooler (ColdEdge, Allentown, USA). The parameters for reduced 2Fe2S and Fe^3+^ citrate measurements were as follows: temperature 20 K, microwave power 2 mW, modulation amplitude 14.36 G, conversion time 40.96 ms, time constant 81.92 ms, sweep time 83.89 s, and the number of scans was 1–3, depending on the signal-to-noise ratio. For the free radical signal, the following parameters differed: temperature 200 K, microwave power 80 mW, modulation amplitude 6 G, and the number of scans was 1–12, depending on the signal-to-noise ratio. All amplitudes of the recorded EPR spectra were normalized using the amplitude of the Fe^3+^ citrate spectra as the internal standard.

## Results

To understand how nature realizes EB at a fundamental level, we conducted QM calculations of possible enzyme-catalyzed reactions using multiple models, with different electron/proton configuration, based on cyt*bc*_1_ crystallographic structures from *R. capsulatus*. For clarity, we specify that in this work, EB is considered as a minimal cascade of internal reactions within the enzyme, leading to a stable, low-energy state containing the product Q, which corresponds to a state that can be detected experimentally as the dominant one under equilibrium conditions, i.e., upon depletion of reaction substrates. In this work, the terms “equilibrium” and “non-equilibrium” are used to refer to experimental conditions in which substrates and products are at redox equilibrium (no net catalytic turnover) and to conditions in which substrates and products have not yet reached equilibrium and the enzyme operates under continuous turnover, respectively. We consider only the primary EB steps, rather than the entire catalytic Q cycle. For the first time, and unlike as in previous studies^53–57^, the QM model of Q_o_ included both acceptors of electrons from QH_2_: 2Fe2S and *b*_L_ (Fig. 1D, E). In the starting state, QH_2_ forms a hydrogen bond to deprotonated H156 of iron-sulfur protein (*R. capsulatus* numbering) and *via* water molecule to E295 of cyt*b*, at the time when immediate electron acceptors 2Fe2S and *b*_L_ are oxidized. Y147 of cyt*b* adopts two conformations: away from or close to E295 (**R** in Fig. 1D or **R**^**Y**^ in Fig. 1E, respectively and Fig. S1). We tested effect of protonation of H156 but in that case no reaction was observed. The choosing deprotonated state of H156 for enzyme-substrate complex was supported by several experimental observations suggesting that protonation of this residue at lower pH inhibits activity of cyt*bc*_1_ ^58–62^. The rationale behind selection of the starting state is explained in SI (Section „Selection of the starting model”).

Initially, following the assumptions of CEB, we tested the scenario in which the reaction is expected to start from **R** or **R**^**Y**^ with protonation of H156 by C4-OH of QH_2_ (atom numbering in Fig. 1F), followed by eT to 2Fe2S. Neither of expected structures containing *b*_L_^ox^, QH^-^, protonated H156 and oxidized 2Fe2S (2Fe2S^ox^), nor those with *b*_L_^ox^, SQH/SQ^-^ and reduced 2Fe2S (2Fe2S^red^) could be obtained in our QM models. Instead, relocation of proton from C4-OH to H156 is accompanied by both: pT from C1-OH to E295 and simultaneous eT from QH_2_ to *b*_L_. This led to the intermediate containing SQ^-^, *b*_L_^red^, and 2Fe2S^ox^ (**I2** in Fig. 2A and **I2** or **I2**^**Y**^ in Fig. S2), which appeared in a clear contradiction with predictions based on CEB, namely SQ formation together with 2Fe2S^red^ and *b*_L_^ox^. Such the state seems quite advanced in the progression of EB as it must have resulted from at least two pTs and one eT (protons are already on H156 and E295 and electron is on *b*_L_). It thus appears that transition from **R**/**R**^**Y**^ to **I2**/**I2**^**Y**^ might involve an intermediate step to be considered. Therefore, we went back to the beginning of the reaction and tested how **R** or **R**^**Y**^ responses to the initial water-mediated pT from C1-OH to E295. Surprisingly, this pT was coupled to eT from QH_2_ to *b*_L_, leading to **I1** or **I1**^**Y**^, (states containing SQH, *b*_L_^red^ and 2Fe2S^ox^, Fig. 2A and Fig. S2). The emergence of *b*_L_^red^ next to 2Fe2S^ox^ was unexpected result, given that *E*_m_s of *b*_L_ and 2Fe2S, determined for proteins under equilibrium, suggests a higher electron affinity for 2Fe2S than *b*_L_. This prompted us to scrutinize the electronic structure of **R**, which was taken as the most probable state of the Q_o_-QH_2_ complex (**R**^**Y**^ and following **I1**^**Y**^ were 1 and 5 kcal/mol less stable than **R** and **I1**, respectively).

**Figure 2.**
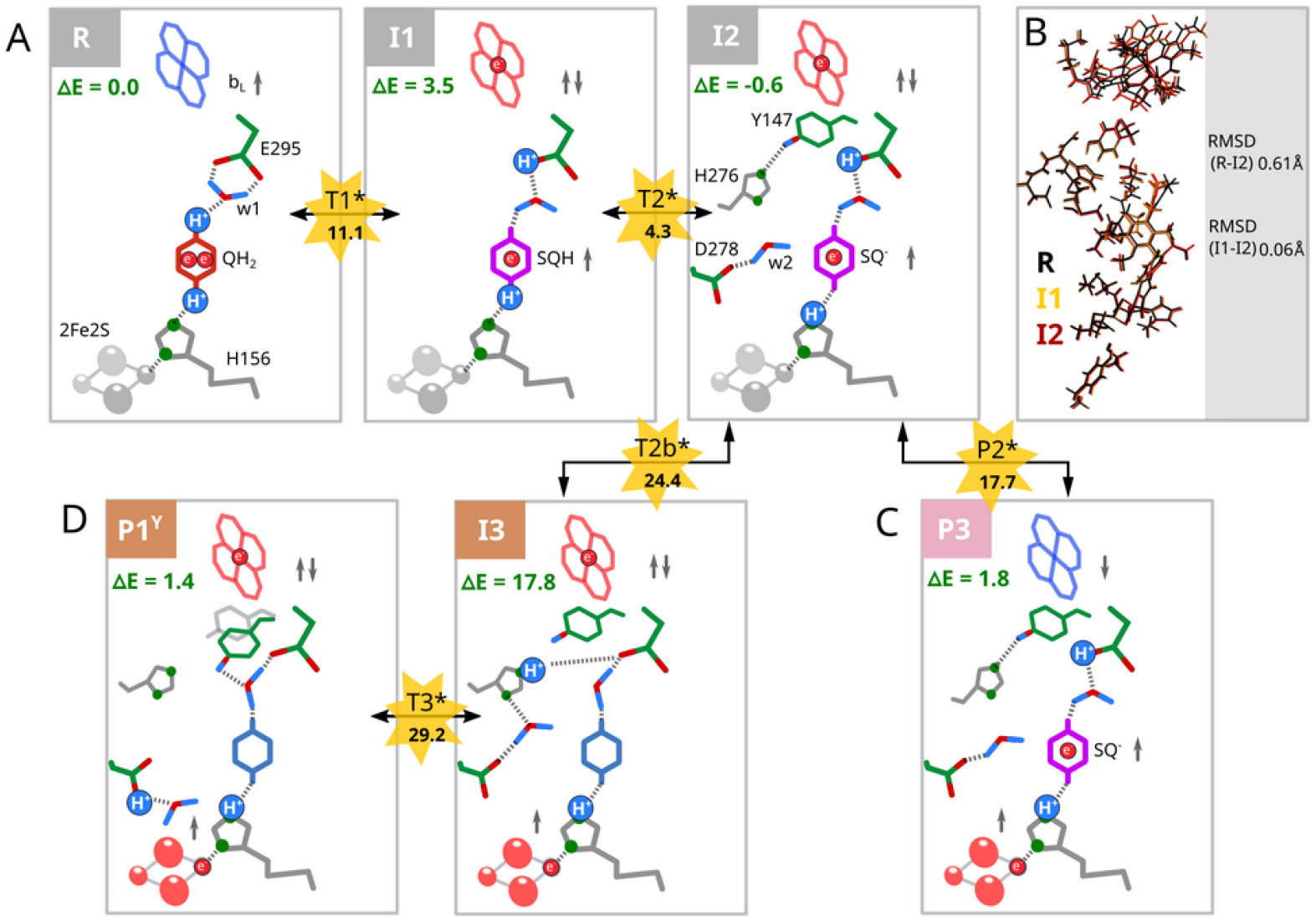
Primary steps of EB without involvement of eT from *b*_L_ to *b*_H_. A) **R** denotes a starting state comprising *b*_L_^ox^ (*blue crossed-hexagon*), QH_2_ (*red hexagon*), and 2Fe2S^ox^ (*grey ovals*). The reaction progresses to **I1** and **I2** states containing *b*_L_^red^ (*red crossed-hexagon*) and SQH, or SQ^-^ (*magenta hexagon*), respectively. B) Superimposed geometries of **R, I1**, and **I2.** C) Evolution of **I2** to stable **P3** containing *b*_L_^ox^, SQ^-^, and 2Fe2S^red^ (*red ovals*), *via* **P2**^**\***^. D) Evolution of **I2** to **P1**^**Y**^ containing *b*_L_^red^, Q, and 2Fe2S^red^ *via* high energy **T2b**^**\***^, **I3** and **T3**^**\***^. Gray arrows indicate the net spin corresponding to the redox state of the substrate and cofactors. Side chains of the protein residues are shown when their protonation and/or conformation state is significant. Important water molecules are labeled as “w1” and “w2” and protons derived from QH_2_ in the given state are represented by blue circles. The energy barriers were estimated *via* energy of pre-TS states (*yellow stars*). Energy is given in kcal/mol.

### Asymmetry in charge and spin distribution in the initial structure

In **R**, there is a shift of electron density from QH_2_ toward *b*_L_, associated with a spread of unpaired α spin density (SD) over M154 and D278 of cyt*b* toward QH_2_ (Fig. 3A). At the same time, α (S_Fe1_=5/2) and β (S_Fe2_ = -5/2) SD are present on iron ions of 2Fe2S^ox^ but antiferromagnetism cancels the net spin. Shifting of electron density toward *b*_L_ imposes a heterogeneous distribution of the local electric potential around the quinone ring with the more negative potential at the region of C1-OH compared to C4-OH (Fig. 3B, red arrow). At the same time, the electronic structure indicates that the highest occupied and the lowest unoccupied molecular orbital (HOMO and LUMO, respectively) are located on QH_2_ and *b*_L_, respectively (Fig. 4). The charge polarization and the location of HOMO and LUMO determines direction of eT from C1-OH to *b*_L_^ox^ accompanied with pT to carboxyl group of E295, resulting in state **I1** (Fig. 2A and S2A). It is important to emphasize that it was not possible to obtain **I1** state when 2Fe2S is not involved (Fig. S3-S6), because in models deprived of the cluster, HOMO orbitals were no longer located on QH_2_ molecule. On the other hand, for the models containing the 2Fe2S cluster but not *b*_L_, we obtained stable state containing SQ-2Fe2S^red^ similarly to previous work^53^ (see optimized geometries and HOMO, LUMO orbitals in Fig. S7). This would indicate that either electron transfer from QH_2_ to 2Fe2S is exoergic, which seems incompatible with CEB, or that it is an artifact of the truncated model. If such a model truly reproduced the first EB step, one would expect to detect an EPR signal of SQ and/or SQ–2Fe2S^red^ under equilibrium conditions and in cyt*bc*_1_ mutant with *b*_L_ knocked out, which is not observed^63^. Clearly, approaches for determining initial steps of EB using models containing only one cofactor at a time suffer from the *a priori* imposition the sequence of the reactions.

**Figure 3.**
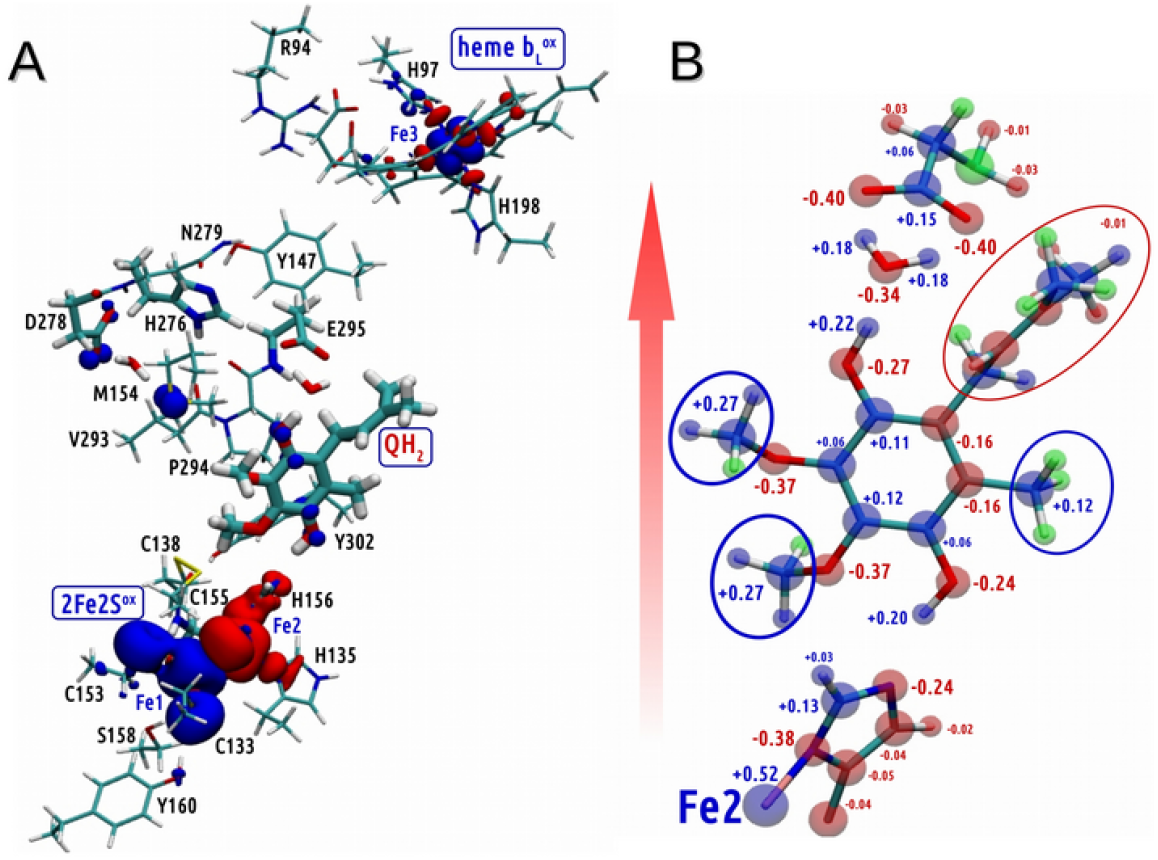
Spin and charge distribution in R. A) The distribution of *α* (blue) and *β* (red) spin density in the most stable structure containing QH_2_. B) The charge distribution within the closest surrounding of the QH_2_ molecule. Red arrow indicates the direction of the shift of negative charge at Q_o_ toward *b*_L_.

**Figure 4.**
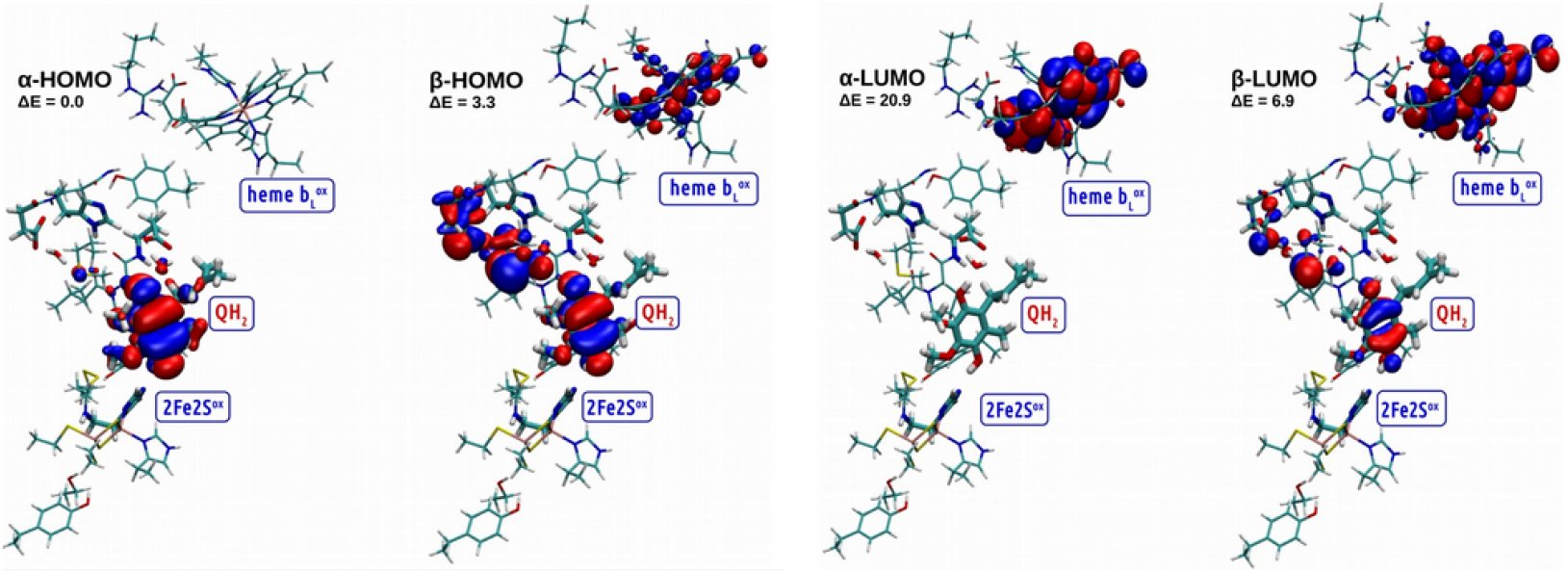
Location of HOMO and LUMO orbitals and their relative energy (with respect to the *α*-HOMO) in kcal/mol for the most stable structure containing QH_2_, *b*_L_^ox^, 2Fe2S^ox^ and deprotonated H156.

Altogether, these observations indicate that reaching of **I1** requires the presence of both *b*_L_ and 2Fe2S, despite apparent lack of involvement of the latter in this process. Furthermore, 2Fe2S and its direct surrounding must be in the correct state, as the models of **R** variants with either protonated H156 and 2Fe2S^ox^ or deprotonated H156 without the cluster also failed to show any signs of progress to **I1** (Fig. S8).

### Reversible EB stages on a way toward product

The state **I1** passes to **I2** by shifting H^+^ originating from the C4-OH group of SQH toward H156. The RMSD parameter comparing the positions of all atoms in **R, I1** and **I2** is about 0.06Å indicating negligible structural rearrangement upon **R↔I1↔I2** transitions (Fig. 2B).

The energy levels of **R, I1** and **I2** are close to each other and the estimated upper-bound barrier for **R↔I1** (11.1 kcal/mol) and **I1↔I2** (0.8 kcal/mol) transitions are relatively small, which means that at room temperature (∼300 K) they are expected to be similarly populated. Thus, **R, I1** and **I2** are considered as degenerate, coexisting states.

We first searched for states containing stable product under conditions when electron on *b*_L_ cannot advance to *b*_H_ (like in antimycin-inhibited enzyme). One tested reaction involved elongation of the H-bond between SQ^-^ and protonated H156. This resulted in unstable **P2**^**\***^ (Fig. 2C and Fig. S9) which immediately relaxed back to **I2** or evolved to **P3**. The appearance of the **P3** state is notable because it contains *b*_L_^ox^ together with SQ and 2Fe2S^red^. Because both of these paramagnetic centers (SQ and 2Fe2S^red^) have parallel spins and are connected with H-bond, one can expect formation of an ferromagnetically coupled SQ-2Fe2S center (discussion in SI, Fig. S10). The state **P2**^**\***^ should be considered as an approximate transition state (pre-TS), with estimated energy +18 kcal/mol, between **I2** and **P3**. We note that nearly identical geometry of nuclei in **I2** and **P3** indicates a potential contribution from electron tunneling process, which effectively decreases the barrier between these states.

The second tested reaction involved shifting the proton away from E295 to D278. This resulted in relatively stable **P1**^**Y**^ containing Q (Fig. 2D). However, as reaching of this state from **I2** is slightly endoergic and required crossing barriers (25 and ∼ 11.4 kcal/mol, Fig. 2) with a high energy state in between (**I3**, +18 kcal/mol, Fig S11), this process is unlikely. Consequently, when electron cannot advance from *b*_L_ to *b*_H_, the enzyme is expected to become trapped in the states **R**↔**I1**↔**I2** and **P3** without possibility to produce a low-energy product Q. This is consistent with inhibitory effect of antimycin on the Q_o_ site.

### Releases from degenerate states and emergence of product

In further search for exoergic processes that would push EB toward low-energy product, we estimated the net energetic cost associated with eT from *b*_L_ through *b*_H_ to Q_i_ (Fig. 5). We considered a state **I2’** which is a result of removal of electron from *b*_L_^red^ in **I2**, and various states of the complementary model that mimicked the acceptance of this electron. The complementary models encompassed *b*_H_ and quinone at Q_i_ (details in SI and Fig. S12). The results indicate that the transition from **I2** to **I2’** would be endoergic (+67,2 kcal/mol) if the complementary model was not included. However, the average energy released per electron delivered to Q_i_ (-95.8 kcal/mol) lowers energy of **I2’** to -28,6 kcal/mol (Fig. 5). In this way, eT from Q_o_ to Q_i_ makes EB exoergic allowing Q_o_ to departure from degenerated states **R**↔**I1**↔**I2** and **P3** (the latter state is unlikely to be formed when this eT is not blocked). The net energy released in Q_i_ drives the enzyme to overcome local barrier associated with pT from E295 to H276 (state **I3’**). The following pT from H276 to D278 is coupled to eT from SQ to 2Fe2S^ox^ which weakens the interaction between quinone moiety and protonated H156 (the H-bond elongates from 1.5 to 2.0 Å, Fig. S13, S14). This reaction is the final step of EB leading to the low-energy state **P1’**^**Y**^ (-23.0 kcal/mol in respect to **R**) containing product (quinone).

**Figure 5.**
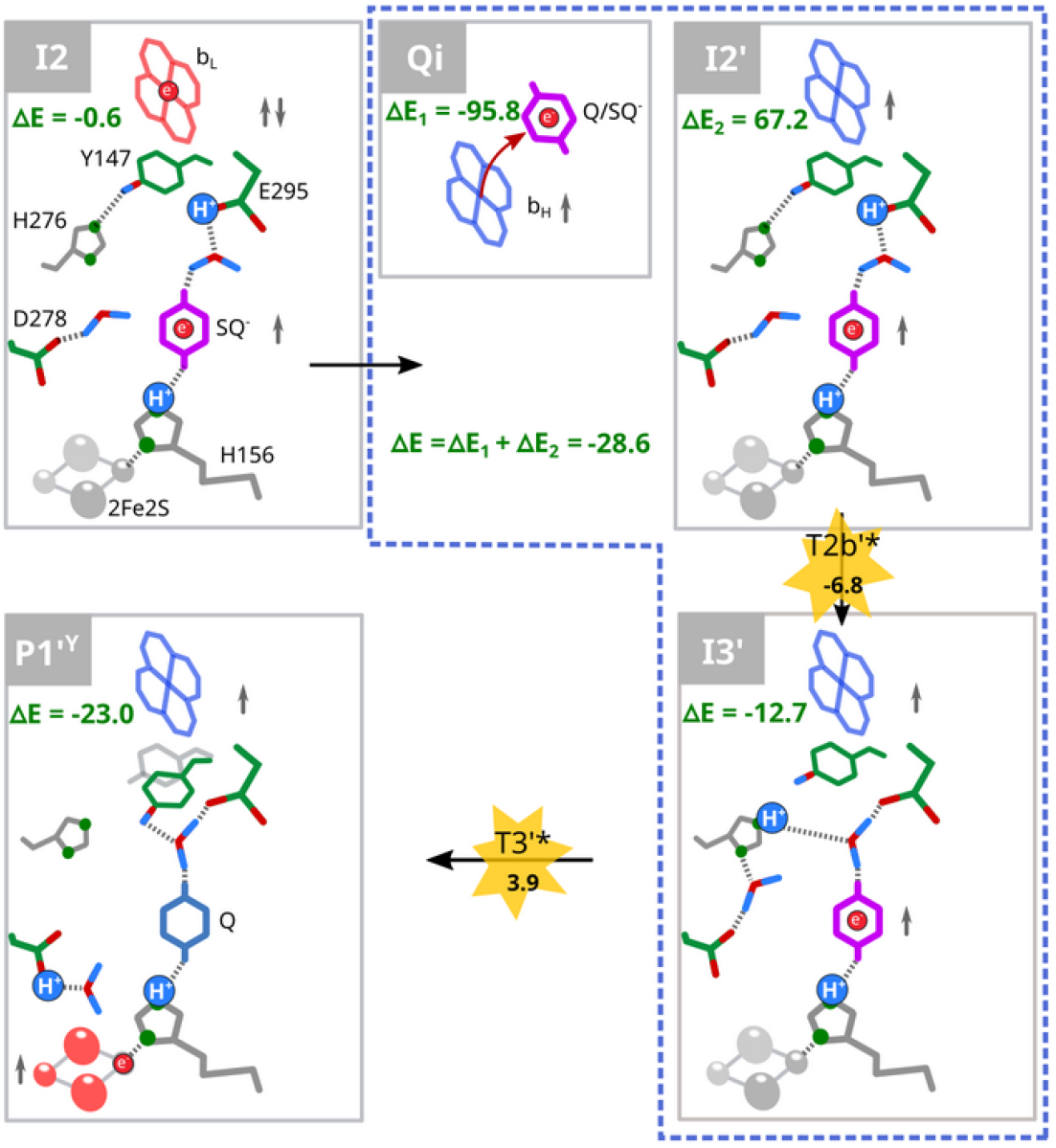
Completion of EB involving energy released during eT from *b*_H_ to Q_i_. The eT from *b*_L_ to *b*_H_ and subsequently to Q_i_ provides a highly exoergic step that resolves the **R↔I1↔I2** degeneration (see Fig. 2), driving the reaction from **I2** toward **I3’.** The energy associated with this reaction was calculated using the states **Q**_**i**_ and **I2’, I3’** considered together (*blue dashed square*). **P1’**^**Y**^ denotes the final state of EB containing Q, 2Fe2S^red^ and protonated D278. The symbols are consistent with those used in Fig. 2. Red arrow indicates eT. The energy barriers were estimated *via* energy of pre-TS states (*yellow stars*). Energy is given in kcal/mol.

We note that while the essence in the departure from degenerated states lies in effective removal of the proton away from the quinone ring, the exact pT path is not a determinant of this process but influences its efficiency. The process might be tolerant to mutational changes and the residues that build the path may vary in different species^64–67^. We also note that, in this mechanism, completion of EB does not require the motion of the ISP away from Q_o_ implicated as inherent feature of catalysis. It it worth mentioning about recent reports on structures of cyt *bc*_1_ for which ISP does not move during the catalytic cycle^68^. When the motion occurs during further steps of enzymatic catalysis that follow **P1’**^**Y**^, transferring electron toward more remote cofactors (*c*-type hemes) it further lowers the energy of the system. However, as mentioned above, these steps remain beyond consideration of work.

Figure 6 summarizes the most probable enzyme states that constitute the pathway of EB-based QH_2_ oxidation inferred from energetic relations between the states obtained by QM (see Fig. S15 for all the considered states). When QH_2_ bound in Q_o_ encounters 2Fe2S^ox^ and *b*_L_^ox^ (**R**), the negative charge from the quinol moiety initially shifts toward *b*_L_ (**I1, I2**). ET from the resulting SQ to 2Fe2S (**I3′, P1′**^**Y**^) is powered by the energy released upon electron transfer from *b*_L_ to Q_i_ (blue line). Without this energetic input, the enzyme is trapped in states **R, I1, I2**, and **P3**, with no possibility of reaching a low-energy product state. These results highlight that productive quinol oxidation cannot be explained by a strictly sequential two-step scheme, but instead requires a cooperative route initiated at *b*_L_^ox^.

**Figure 6.**
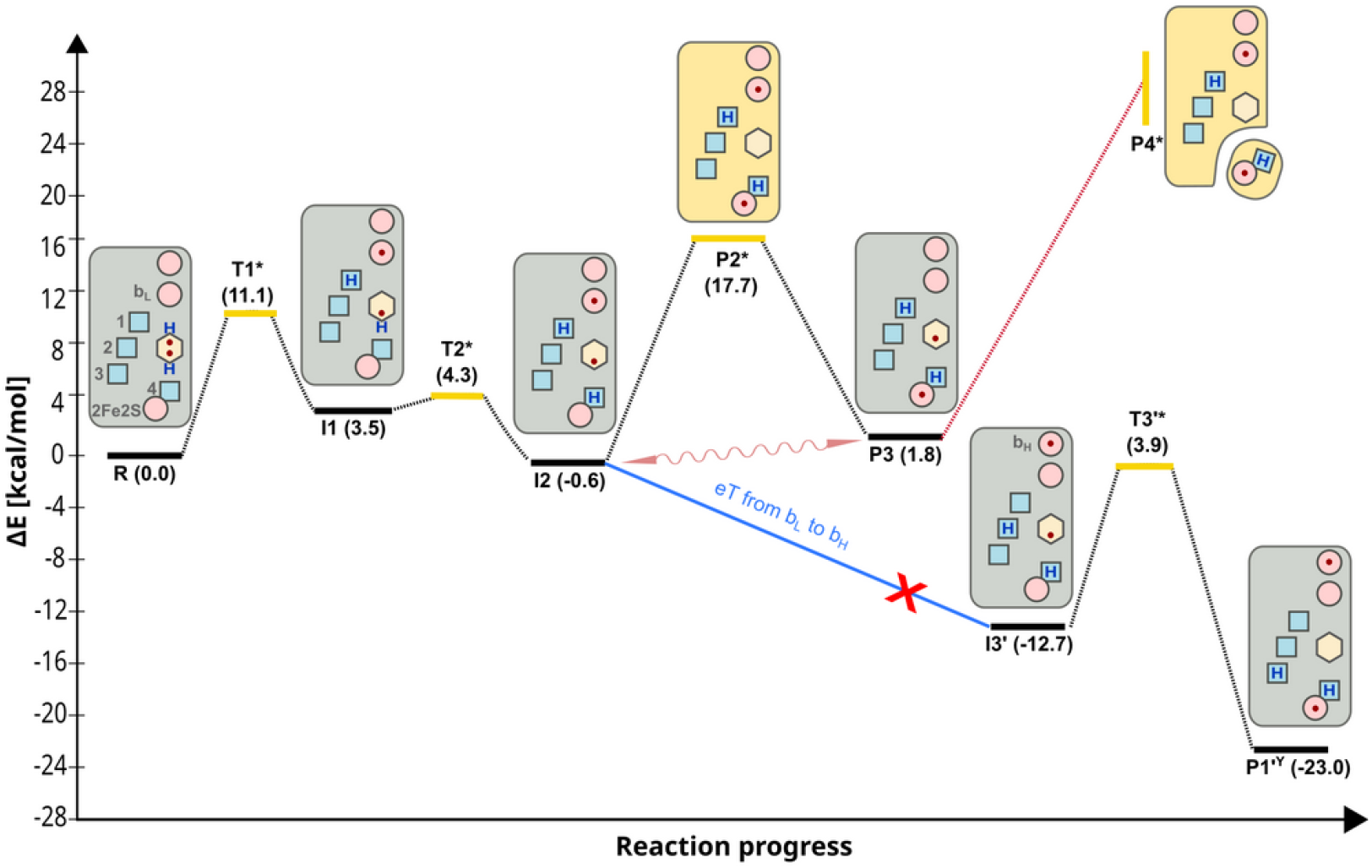
Key states of the EMBER model of EB. Solid black lines represent relative energy levels (numbers in parentheses) of the states on the way from the substrate (**R**) to product (**P1’**^**Y**^). Yellow solid line represents energy level of pre-TSs (**T1**^**\***^, **T2**^**\***^, **T3**^**\***^, **P2**^**\***^). Red circles – electron acceptors; blue squares – proton acceptors 1, 2, 3 in cyt *b* (in *R. capsulatus* E295, H276, D278, respectively), and 4 in ISP (histidine ligand to 2Fe2S); yellow hexagon – quinone moiety; red dots and blue “H” – electrons and protons derived from QH_2_; red cross - inhibition by antimycin; wavy arrow – electron tunneling; dotted lines illustrate barriers between states. Red dotted line denotes the possibility of quinone formation **P4**^**\***^ from **P3** by elongation of the distance between ISP and the Q_o_. The energy level of **P4**^**\***^ increases due to uncompensated breaking interactions, thus cannot be determined precisely (yellow vertical line).

## Discussion

In this study, we performed QM calculations to explore possible mechanism underlying the initial stages of electron bifurcation (EB). Our motivation arose from the interpretative challenges posed by experimentally observed enzyme states, particularly under antimycin inhibition. The models explicitly included both immediate electron acceptors (2Fe2S^ox^ and *b*_L_^ox^) and proton acceptors at each reaction step. This approach avoided the predefined sequence of events inherent to the CEB models, in which the reaction is partitioned into two sequential one-electron transfers: first from QH_2_ to 2Fe2S^ox^, then from SQ to *b*_L_^ox^. Optimized stationary points, together with EPR spectroscopic evidence, revealed an alternative to the canonical CEB mechanism, in which EB is initiated by electron transfer from QH_2_ to *b*_L_^ox^ and proceeds along an energetically favorable cooperative pathway with engagement of the integrated system of redox cofactors. To distinguish this mechanism from CEB, we introduce the term EMBER (EMergent B_L_-first Electron Route), a conceptual model that emphasizes the alternative EB pathway not anticipated within conventional CEB assumptions.

An important aspect of the EMBER mechanism is that the initial shift of negative charge toward *b*_L_^ox^ does not occur in models of **R** deprived of 2Fe2S (Fig. S3–S6). In this case, although the LUMO orbitals are still present on *b*_L_^ox^, the HOMO orbitals do not localize on the QH_2_ molecule. This result automatically precludes the eT process in such a model. Additionally, this also eliminates the possibility of reproducing the sequence of reactions identified in our *ab initio* modeling by splitting it into two separate one-electron reactions, since in that case we would, *de facto*, be calculating an irrelevant system. Nevertheless, we checked whether the sequence of reactions predicted by CEB-based models could be reproduced by analogous splitting into two reactions. In this case, we tested the possibility of the first eT from QH_2_ to 2Fe2S^ox^ in a model of **R** deprived of *b*_L_. We found that SQ along with 2Fe2S^red^ were formed but, contrary to the assumptions of CEB, this the state has lower energy than **R**. This means that the process turned out to be exoergic, leading to the formation of stable SQ at Q_o_. In contrast, the complementary reaction of the second eT from SQ to *b*_L_ ^ox^ (reproduced in **P3** deprived of 2Fe2S) was found to be endoergic. This introduces a high energy state that cannot be easily reached. As the electron does not reach *b*_L_, further eT toward Q_i_ is not possible and the system is expected to become trapped in a state with relatively stable SQ, potentially detectable by EPR under equilibrium conditions. This outcome is inconsistent with both CEB assumptions and experimental observations. Thus, splitting EB into two separate eT reactions appears to introduce artifacts, either preventing the initial reaction or producing a stable SQ. Therefore, the occurrence of EB must be viewed as an inherent emergent property of QH_2_ oxidation at Q_o_.

### Predictions inferred from the EMBER model

Our results revealed a fundamentally new picture of the primary steps of EB catalyzed by cyt*bc*_1_, substantially different from CEB. While it is virtually not possible to point to a single experimental observation that would directly validate this mechanism, we highlight several key predictions of the EMBER model along with corresponding experimental observations that, taken together, support the proposed mechanism.

#### Prediction 1

According to the EMBER model, antimycin-inhibited cyt*bc*_1_ exposed to substrates becomes trapped in the states **R**↔**I1**↔**I2**↔**P3** (Fig 6). Among these states, **P3**, containing SQ^-^ and 2Fe2S^red^ with parallel spins, appears to be the most characteristic spectroscopically detectable species. The structural proximity of these two centers, connected by an H-bond provides a strong argument for the origin of the spin-coupled SQ-2Fe2S^red^ EPR signal^18,19,69^. This signal exhibits a rhombic spectrum arising from spin-spin exchange interaction with estimated frequency ∼3.6 GHz, producing the most prominent transition at g = 1.94 (Fig. 7A, green) - clearly distinguishable from the 2Fe2S^red^ signal at g = 1.90 (Fig. 7A, black). It can be further suggested that the amplitude of SQ-2Fe2S^red^ signal should correlate with the amplitude of the SQ radical signal, posed by **I1** and/or **I2** if the system is in the stationary state during the turnover. Therefore, one would expect the SQ signal to be detectable by EPR in the same samples containing SQ-2Fe2S^red^. Indeed, several reports describe distinct SQ signals under non-equilibrium conditions in antimycin-inhibited cyt*bc*_1_ including ours^18–22,70,71^. In this work, we also observed SQ signal at temperatures above 150 K, together with SQ-2Fe2S^red^ at 20 K (Fig. 7B). Interestingly, these two signals were found to be linearly correlated, in agreement with expectations derived form the EMBER model. While the SQ–2Fe2S^red^ signal provides a specific signature of spin coupling between two defined cofactors, assigning the SQ signal itself to **I1** and/or **I2** remains inherently challenging. At this stage, we cannot rule out a possibility that the detected SQ arises from a thermally activated process related to the SQ–2Fe2S^red^ state.

**Figure 7.**
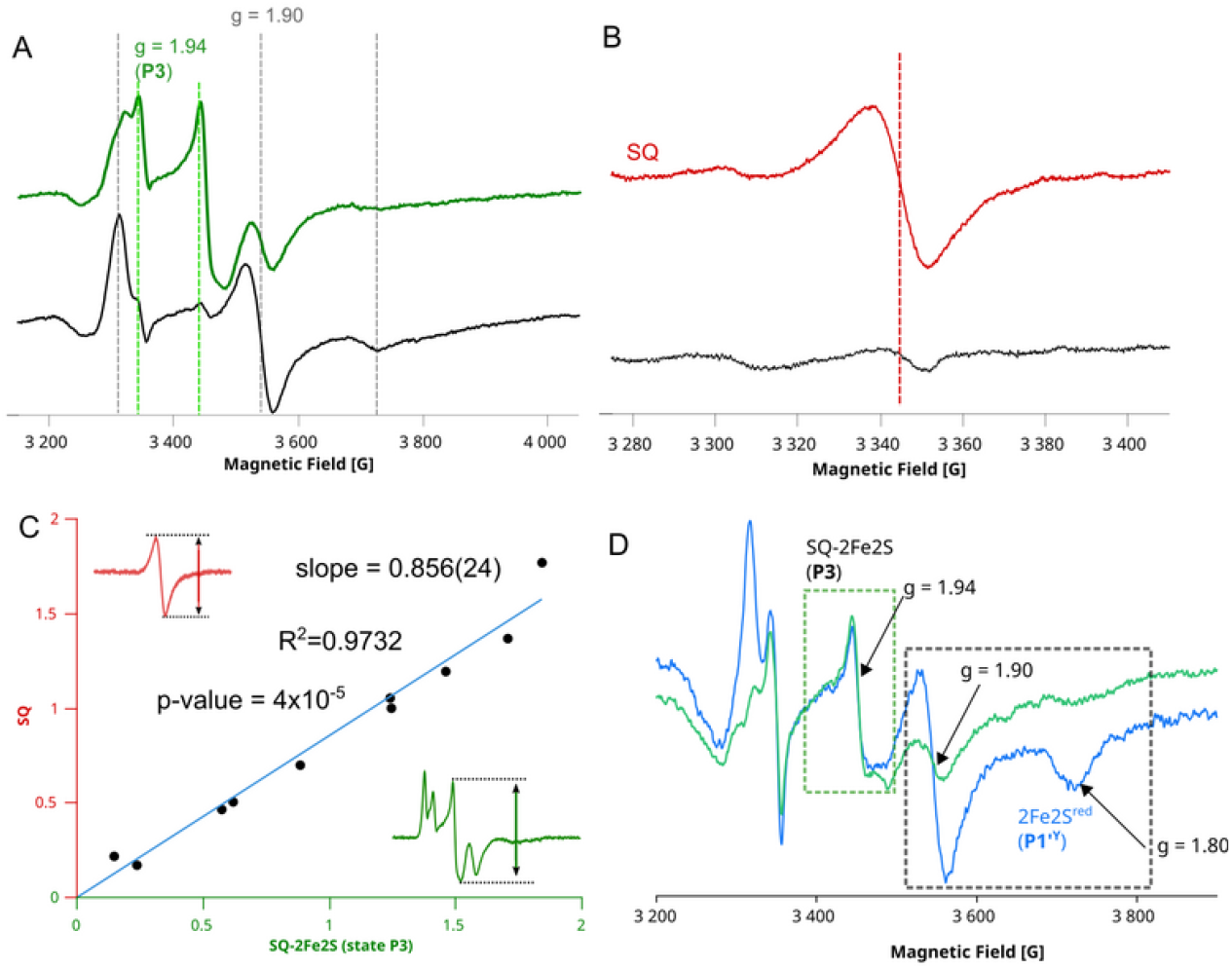
Experimental data obtained for antimycin-inhibited cyt*bc*_1_. (A) EPR spectra of 2Fe2S^red^ and coupled SQ– 2Fe2S^red^ (**P3**) measured at 20 K for samples prepared under equilibrium (black) and non-equilibrium conditions (green; 1 s after substrate addition). Dashed vertical lines indicate the main EPR transitions of 2Fe2S^red^ (gray) and SQ–2Fe2S^red^ (green). (B) SQ radical signal measured at 200 K in the same samples as in A, under equilibrium (black) and non-equilibrium (red) conditions. (C) Correlation between the amplitudes of SQ and g = 1.94 EPR signals for samples prepared at different time points after mixing with substrates. (D) Double-mixing EPR experiment showing spectra of enzyme pre-set to **P3**, followed by mixing with buffer (*blue*) or with an additional portion of oxidized cyt*c* (*green*). Grey and green dashed boxes highlight spectral ranges corresponding to 2Fe2S^red^ with bound Q and SQ–2Fe2S^red^ (**P3**), respectively.

#### Prediction 2

When the reaction reaches equilibrium after substrate exhaustion, both the SQ and SQ–2Fe2S^red^ EPR signals disappear (Fig. 5A,B; red and black, respectively). This provides clear evidence that they originate from species formed exclusively under non-equilibrium conditions and therefore represent short-lived, experimentally detectable states that are populated only under continuous turnover conditions. Moreover, EMBER predicts that **P3** occurs in enzyme molecules in which *b*_L_ remains oxidized. Accordingly, the signal at g = 1.94 should be accompanied by the *b*_L_ ^ox^ signal, a result consistently reproduced in experiments^18,19^.

#### Prediction 3

The transitions between **I2** and **P3** are relatively slow as they require crossing **P2**^***\****^, which lies ∼18 kcal/mol above both **I2** and **P3**. This barrier corresponds to the rate constant of approximately 1/s ^72^. The experimentally obtained rate of growth of SQ-2Fe2S^red^ signal (Fig. 8 green) is ∼4.7/s, which, according to Eyring equation, corresponds to the energy barrier ∼16 kcal/mol. If the observed growth rate of the SQ–2Fe2S^red^ signal indeed reflects the internal barrier-crossing step rather than substrate diffusion, then the close match with the calculated values, within this assumption, supports the reliability of the energy profile in Fig. 6.

**Figure 8.**
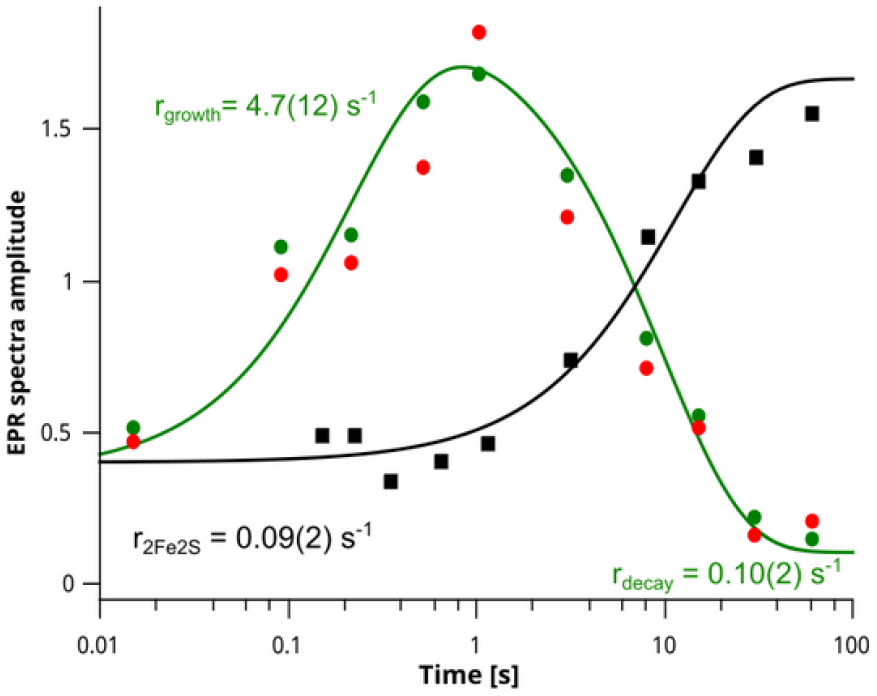
Time dependence of EPR signals in antimycin-inhibited cyt*bc*_1_. Green and red circles represent the amplitudes of SQ–2Fe2S^red^ (g = 1.94) and SQ (g = 2.0), respectively. Black squares denote the amplitude of 2Fe2S ^red^ with Q bound at the Qo site. The solid green line corresponds to a biexponential fit (growth and decay) of the SQ– 2Fe2S^red^ signal, while the solid black line represents a single-exponential fit of the 2Fe2S^red^ signal.

We further propose that a tunneling process (eT exchange between 2Fe2S, Q and *b*_L_) may contribute to the **I2** ↔ **P3** transitions, effectively lowering this barrier.

The observed decrease of SQ-2Fe2S^red^ at the rate ∼0.1/s is explained by a spontaneous *b*_H_ ^red^ oxidation. It reopens possibility of eT from *b*_L_ to *b*_H_ eT and progression toward the final **P1’**^**Y**^. As a consequence, one may observe decrease in amplitude of 2Fe2S^red^ with Q formation at Q_o_ at the same rate as SQ-2Fe2S^red^ decay (Fig. 8 black). This rate is comparable to antimycin-inhibited turnover of cyt*bc*_1_.

#### Prediction 4

The relatively slow predicted kinetics of **P3** formation, together with the parallel spins in SQ and 2Fe2S^red^, provide strong arguments against the possibility that SQ– 2Fe2S^red^ arises as the initial step of EB resulting from electron transfer from QH _2_ to 2Fe2S^ox^. In particular, further transition of **P3** *via* the high-energy **P4**^**\***^ intermediate would yield a product at Q_o_ and complete EB. According to CEB, such a process would entail the risk of short-circuit reactions during subsequent QH_2_ oxidation to SQ, because *b*_L_ (remaining reduced from the previous cycle) could rapidly re-reduce SQ back to QH_2_. To avoid such short circuits, the enzyme must minimize this pathway, for instance by making the **P3** → **P4**^**\***^ transition highly unfavorable. In Fig. 7D, we show the effect of blocking 2Fe2S^red^ oxidation in **P3**. When antimycin-inhibited cyt*bc*_1_ was mixed with a sub-stoichiometric amount of substrates, the system reached a stationary state (that can be ascribed to **P3)** after 1 s. At this stage, the oxidizing capacity (oxidized cyt*c*) was nearly exhausted. Under these conditions, a fraction of the enzyme contained SQ–2Fe2S^red^ (Fig. 7D, blue spectrum, g = 1.94), while another fraction contained reduced 2Fe2S^red^ with Q bound at Q_o_ (Fig. 7D, blue spectrum, g = 1.90). The Q molecule bound at the Q_o_ site is evidenced by the g = 1.80 characteristic transition^73^. Upon mixing the sample with an extra portion of cyt*c* (increasing oxidizing power), the 2Fe2S^red^ signal decreased dramatically upon oxidation (Fig. 7D, green spectrum), whereas the signal corresponding to **P3** remained unchanged. This provides strong suggestion that in the **P3** state, 2Fe2S^red^ cannot leave Q_o_ to generate Q, and thus the transition from **P3** to **P4**^**\***^ is not possible.

#### Prediction 5

The existence of **P3** resembles a “dead-end” branch in the QH_2_ oxidation pathway, blocking Q_o_ and reducing enzymatic activity from QH_2_ oxidation pathway, blocks Q_o_ and reduces the enzymatic activity from ∼400/s to approximately 4/s ^74^. This blocking effect may explain why antimycin inhibits cyt*c* reduction by cyt*bc*_1_. In the absence of the inhibitor, when *b*_H_ can accept an electron from *b*_L_, **I1** and **I2** remain transient, whereas **P3**, being kinetically isolated, does not accumulate to significant levels. Consequently, the paramagnetic signatures of **I1, I2**, and **P3** should remain undetectable by EPR under equilibrium conditions or in the non-inhibited enzyme. This expectation has been confirmed by previous experiments^75^.

#### Prediction 6

Formation of **P3** requires prior eT to *b*_L_ (i.e., transition from **R** to **I1**/**I2**). Therefore, a mutation that disrupts the normal redox activity of *b*_L_ should eliminate **P3**. Indeed, the spectroscopic signature of **P3** was not detected in a mutant with impaired *b*_L_ function, caused by substitution of one of its axial histidine ligands with asparagine^63^.

#### Prediction 7

It can be proposed that the energy released during eT from *b*_L_ to Q_i_ facilitates the movement of ISP, thereby enhancing the probability of rapid eT from 2Fe2S^red^ to cyt*c*_1_. Indeed, although indirect, experimental evidence supporting the existence of such a mechanism has been reported^76^.

### Large models *vs* transition states estimation

Several attempts have been made to calculate TSs states in modeling EB; however, these models were limited only to one of the active redox cofactors. Thus, they usually included the QH_2_ and 2Fe2S^ox 53–57^ or SQ and heme *b*_L_^ox^ at a time imposing the reaction following assumption of CEB^53^. We employed large cluster models (∼400 atoms) because dividing the system into smaller fragments centered on individual cofactors led to qualitatively different electronic states, resulting in a biased energetic landscape (SI, Figures S3–S8). However, the use of such large QM models renders full transition-state characterization impractical due to the prohibitive cost and numerical challenges associated with evaluating the full Hessian matrix (∼1200 × 1200 in the present case) ^*38*^ . Consequently, constrained TS-like structures (pre-TS) were used to obtain conservative upper-bound estimates of the activation barriers. In standard transition-state search protocols, pre-TS geometries serve as initial guesses for TS optimization, which relies on vibrational analysis to locate a first-order saddle point on the potential energy surface; full optimization typically yields a relaxed TS structure at a lower energy than the initial pre-TS geometry.

### Equilibrium redox potentials *vs* state-dependent electron affinities of cofactors engaged in EB

According to EMBER, the catalytic reaction is initiated by concerted pT from QH _2_ to E295 and eT from QH_2_ to *b*_L._. This reaction has never been considered by CEB, as in the framework of equilibrium electrochemical theory, the *E*_m_ of *b*_L_ appears too low for an effective withdrawal of electron from QH_2_. Reconciliation of this apparent contradiction lies in a postulation that values of redox potentials of cofactors determined under equilibrium might not be relevant when describing individual each step of the reaction, but rather reflect the final thermodynamic relationships between the reduced and oxidized populations of all redox elements once the system has reached equilibrium and the reaction is complete. While this statement may seem controversial, it does not challenge the general validity of Marcus theory^77^. Rather, it allows us to consider that under certain transient conditions present in cyt*bc*_1_ (and possibly in other multicofactor enzymes), additional dynamic factors, such as charge separations due to eT and pT, may effectively modulate electron affinity around particular redox cofactors fixed within a protein environment of relatively low dielectric constant. This modulation may arise from dynamic changes in the protonation states of amino acid residues and in the redox states of interacting cofactors during the course of the reaction. Indeed, previous studies have shown that reduction of one heme can lower the redox potential of an adjacent heme through Coulombic interactions by tens or even hundreds of millivolts^78^. Such changes in electrostatic interactions often lead to splitting or stretching of Nernst curves (due to a lowering of the n parameter below 1) measured for cofactors spatially constrained by the protein structure^79,80^. The effect of such interactions has also been demonstrated for electron distribution in dimeric cyt*b*_6_*f* ^81^. In the case of hemes *b* at distances similar to those in cyt*bc*_1_, shifts of up to ±100 mV should be expected^27,78^. Moreover, protonation of H156 near 2Fe2S can shift the cluster potential from approximately -120 mV to +300 mV ^82^. The latter two examples have direct relevance to description of transient conditions of EB.

Considering that determined *E*_m_ for *b*_L_ corresponds to conditions where *b*_H_ is already reduced, and for 2Fe2S^red^ where protonation of H156 has already occurred, these values clearly do not reflect the state of the activated enzyme–substrate complex in which EB begins. At the onset of EB, the electron affinities of *b*_L_^ox^ and 2Fe2S^ox^ may be relatively similar from the ‘QH2 point of view,’ because at this stage *b*_H_ is oxidized and H156 at the 2Fe2S cluster is deprotonated (protonation prevents the reaction). This effect likely elevates the electron affinity of *b*_L_, lowering the energy levels of **I1** and **I2**. Consequently, a large potential separation between Q/SQ and SQ/QH_2_ is not required at the early stages of EB. After bifurcation is completed and equilibrium is established, the system adopts a state in which *b*_L_ and 2Fe2S exhibit potentials of approximately – 120 mV and +300 mV, respectively, as typically determined from equilibrium redox titrations^83^. CEB, using these values to describe the initial stages would require introducing a large split in the redox potentials of the Q/SQ/QH_2_ triad and, associated with this assumption, alternating sequence of uphill and downhill steps. Consequently, CEB faces a mechanistic issue of having to put additional constraints that would explain prevention of potential quinone-mediated short circuiting of the system. EMBER, proposing a relatively flat energy profile for EB without alternating uphill/downhill steps, is devoid of this concern.

The EMBER model presented here considers the EB mechanism catalyzed by cyt *bc*_1_ at a fundamental level, providing a comprehensive explanatory framework for both current and future experiments. We believe that applying this model and moving beyond the canonical electron bifurcation concept allows several previously difficult-to-interpret observations to be clarified, and those that appeared contradictory can now be unified. As the proposed, concept of transient changes in electron affinities may apply not only to cyt*bc*_1_ but also to other multicofactor enzymes, future experiments should aim to confirm or falsify this model.

## Data availability statement

The experimental data supporting the findings of this study have been deposited in the RODBUK Cracow Open Research Data Repository under DOI: https://doi.org/10.57903/UJ/LVJICQ The data are currently under embargo and will be made publicly available upon publication of this article.

## Authors contributions

Conceptualization: AWA, ŁB, AO, MS. Methodology: AWA, ŁB, MS. Investigation: AWA, ŁB, MS. Visualization: AWA, MS. Funding acquisition: MS. Supervision: MS. Writing – original draft: AWA, MS. Writing – review & editing: AWA, ŁB, AO, MS.

