## Supplementary Information for "Electron bifurcation arises from emergent features of multicofactor enzymes"

**The PDF file includes:**

Supplementary Text

Figs. S1 to S15

**Supplementary text**

Selection of the starting model

To define the starting point for EB reaction, we initially optimized two structures (**R** and **R1**), which contain $QH_2$ and $b_L^{ox}$ and 2Fe2S$^{ox}$ (Fig. S1). In the structure **R1**, $QH_2$ interacts directly with cyt*b*:E295 through the -OH group at C1 position of $QH_2$ (C1-OH), without involvement of water, whereas in **R**, this interaction is mediated by water molecule. The structure containing the water molecule between E295 and $QH_2$ (**R**) is 15 kcal/mol (651 meV) more stable than structure **R1**. This stability difference is associated with the strengthening of the hydrogen bond between the C4-OH group of $QH_2$ and the $N_\tau$ atom of ISP:H156 (ligand to 2Fe2S), as evidenced by the shortening of the distance between the H and $N_\tau$ atoms from 2.0 Å in **R1** to 1.8 Å in **R**. Additionally, the water molecule also contributes to the formation of a hydrogen bond network linking $QH_2$, E295, cyt*b*:H276, and cyt*b*:D278, facilitating proton transfer (pT) from $QH_2$ to bulk water with minimal geometric rearrangement. In general, all states derived from **R** were more stable than **R1**, so we focused only on the structures where water-mediated $QH_2$ binding to E295 served as the starting point for EB (**R** and **R**$^Y$ in Fig. S1, A).

Since cyt*b*:Y147 appears to be involved in proton transfer away from $QH_2$, we also optimized structures containing $QH_2$ in which the tyrosine side chain is rotated to interact with the quinone ring (**R**$^Y$) (Fig. S1, B). All structures with this conformation, where the Y147 side chain is rotated towards the quinone ring, are labeled with the "Y" superscript. Because **R**$^Y$ is only negligibly less stable (1 kcal/mol) than **R**, both structures were considered as possible initial states for EB to start from.

Relation between **I2**, **P2*** and **P3**

**P2*** containing $b_L^{red}$, Q, 2Fe2S$^{red}$ (Fig. 2C and Fig. S9) forms upon stretching of the hydrogen bond between SQ$^-$ and H156. This state could be optimized after imposing constraints on the distance between the C4 carbonyl oxygen of Q and the proton associated with the $N_\tau$ nitrogen of H156. When the distance exceeded 3.0 Å, spontaneous electron transfer (eT) from SQ$^-$ to 2Fe2S$^{ox}$ occurred. **P2*** can be considered an approximation of the barrier between **I2** and **P3**. The latter state forms from **P2*** by transferring the α electron from $b_L$ to 2Fe2S and is only 1.8 and 2.4 kcal/mol less stable than **R** and **I2**, respectively. The **P3** state contains $b_L^{ox}$ and SQ$^-$ ferromagnetically coupled to 2Fe2S$^{red}$, which can be detected by EPR spectroscopy. Alternatively, when the β electron is transferred instead of the α electron, the antiferromagnetic coupling inevitably leads to the oxidation of 2Fe2S$^{red}$ and the reformation of **R** (Fig. S10).

Complementary model mimicking eT from $b_L$ to the $Q_i$ site

The energy diagrams in Fig. 6 and S15 were constructed following the approach proposed by Per Siegbahn and Margareta Blomberg [1] and represent the final energies of the most stable stationary points of the reaction associated with electron transfer (eT) from $b_L$ to $b_H$. These energies were corrected for the energy costs of Q reduction at the $Q_i$ site. The final values were estimated using electronic energies, computed at the same theoretical level for the reduction of Q at the $Q_i$ site and the oxidation of $b_L$.

To include the energy associated with eT from $b_L$ to the $Q_i$ site through $b_H$, additional calculations (using the same computational methods) were performed for the model involving the $Q_i$ site. The model was composed of $b_H^{ox}$, and thirteen amino acid residues around the heme and the $Q_i$ site, and bound Q (the state $Qi^{ox}$ in Fig. S12A). Adding one electron to the state $\mathbf{Qi^{ox}}$ (mimicking eT from $b_L$) leads to spontaneous transfer of this electron to Q, forming $SQ^-$ (state $\mathbf{Qi^{red1}}$). The $SQ^-$ can be protonated to form SQH (state $\mathbf{Qi2^{red1}}$), which slightly stabilizes the semiquinone at the $Q_i$ site. Adding the second electron (mimicking next eT from $b_L$) leads to formation of either SQH and $b_H^{red}$ (state $\mathbf{Qi^{red2}}$) or $QH_2$ and $b_H^{ox}$ (state $\mathbf{Qi2^{red2}}$) with the latter being 8.7 kcal/mol more stable (Fig. S12). $\mathbf{Qi^{red1}}$ is by 108.5 kcal/mol more stable than $\mathbf{Qi^{ox}}$ (ΔE'), while $\mathbf{Qi^{red2}}$ is by 83.2 kcal/mol more stable than $\mathbf{Qi^{red1}}$ (ΔE"). Thus, we estimate the average energy ($\Delta E_1$) released at the $Q_i$ site after accepting one electron from $b_L$ is 95.8 kcal/mol on average ($\Delta E_1 = 1/2\Delta E' + 1/2\Delta E''$). The previously obtained results revealed that pT transfer reaction within the $Q_i$ site runs spontaneously or through negligible barrier [2]. We hypothesize that energy released at $Q_i$ is used to overcome barrier of the subsequent steps of pT and eT at the $Q_o$ site. This hypothesis is consistent with experimental data, which revealed that switching off the $Q_i$ active site significantly affects the $Q_o$ reactions and leads to **P3** formation, which is not an intermediate of the EB in non-inhibited enzyme.

To study the effect of eT from $b_L$ toward $Q_i$ on the further EB process at the $Q_o$ site, we applied the optimized structures **I2'**, **I3'** and **P1'**, which were obtained by removing the electron from $b_L$ from the structures **I2**, **I3** and **P3**, respectively. The energy of **I2'**, **I3'** and **P1'** states were estimated by accounting the energy released by eT from $b_L$ to the $Q_i$ site.

### Supplementary figures

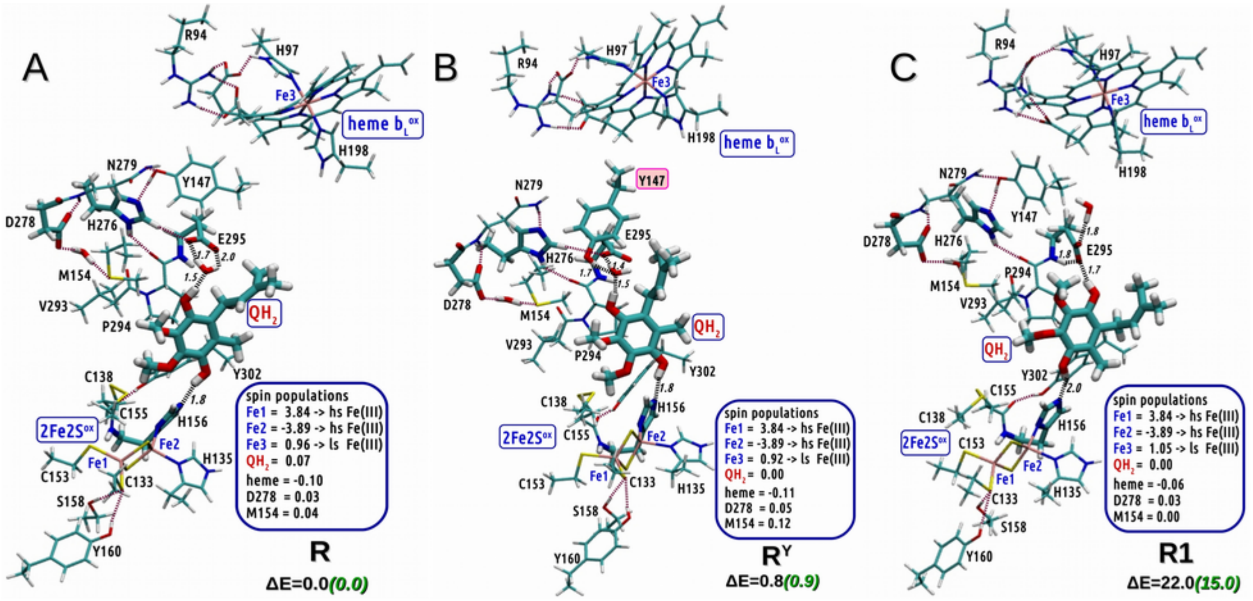


**Figure S1**: The three optimized structures of $Q_o$ sites containing $QH_2$, considered as potential starting points for EB. A) In **R**, $QH_2$ interacts with E295 *via* water molecule. B). In **R**$^{Y}$, $QH_2$ interacts with E295 and Y147 *via* water molecule, C) **R1**, $QH_2$ directly interacts with E295. The relative energy (ΔE) for each of the structure is defined as kcal/mol differences from structure **R**. Black numbers in ΔE show relative energy computed with def2-SVP basis set in vacuum, while green numbers show respective energy using def2-TZVP basis set in combination with PCM modeling protein environment defined by dielectric constant of 4 and radius probe of 1.4 Å. Data shown in blue frames represent spin populations gathered for the important fragments. Distances larger than 2.5 Å are not shown.

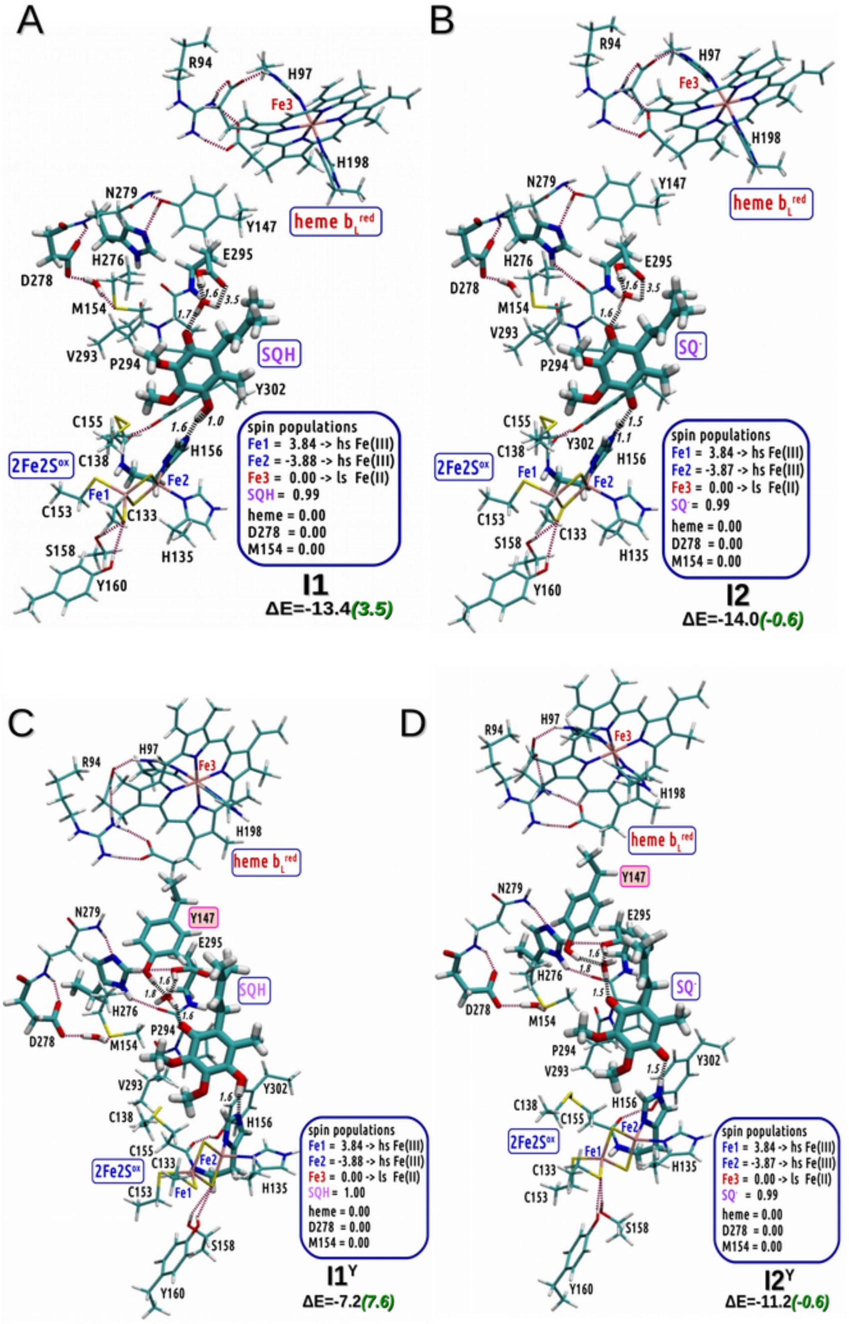


**Figure S2:** The most stable optimized structures containing SQH and $SQ^-$ at the $Q_o$ site. A) The state **I1** containing $b_L^{red}$, SQH and $2Fe2S^{ox}$. B) The state **I2** containing $b_L^{red}$, $SQ^-$, $2Fe2S^{ox}$ . C) The state $\mathbf{I1^Y}$ containing $b_L^{red}$, SQH, $2Fe2S^{ox}$ with the side chain of Y147 rotated toward SQH. D) The state $\mathbf{I2^Y}$ containing $b_L^{red}$, $SQ^-$, $2Fe2S^{ox}$ with the side chain of Y147 rotated toward $SQ^-$. The relative energy (ΔE) for each of the structure is defined as kcal/mol differences from structure **R**. Black numbers in ΔE show relative energy computed with def2-SVP basis set in vacuum, while green numbers show respective energy using def2-TZVP basis set in combination with PCM modeling protein environment defined by dielectric constant of 4 and radius probe of 1.4 Å. Data shown in blue frames represent spin populations gathered for the important fragments. Distances larger than 2.5 Å are not shown.

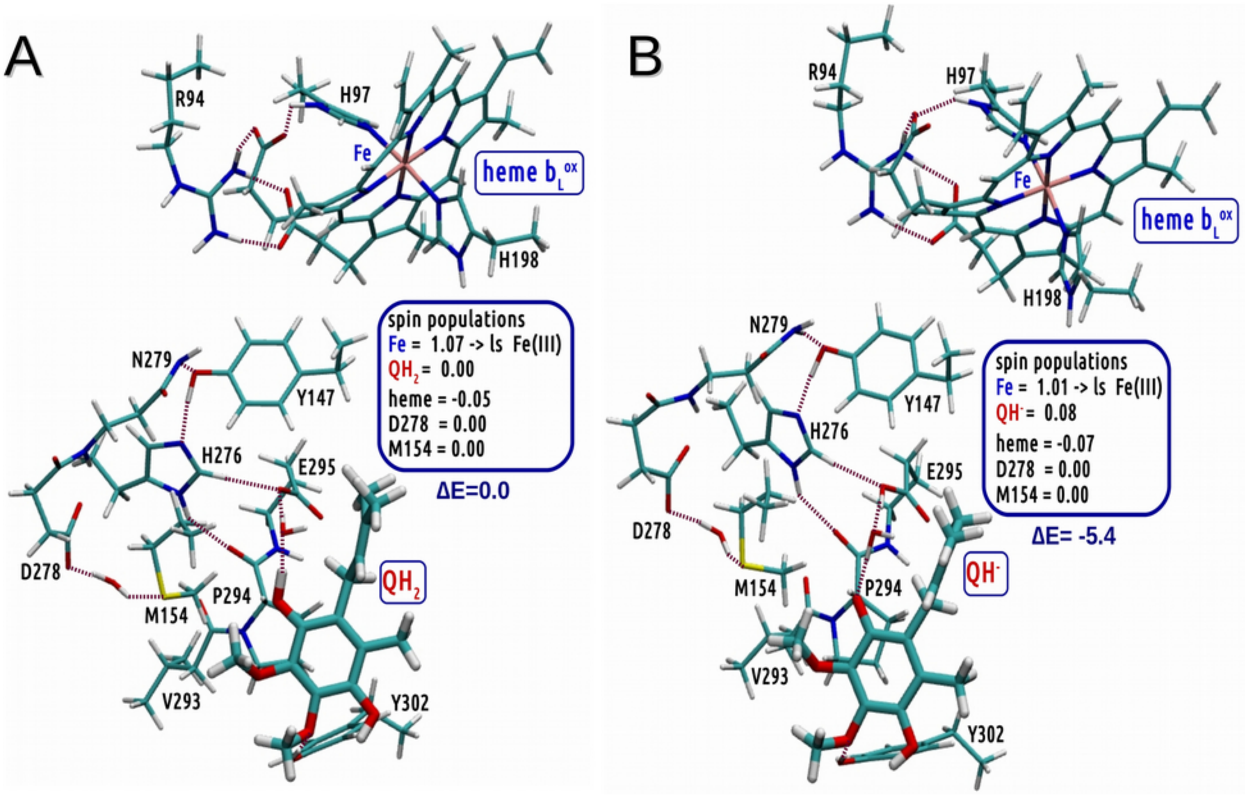


**Figure S3**: Optimized structures containing substrate and $b_L^{ox}$ but without the part housing 2Fe2S. A) The structure representing $QH_2$. B) The structure containing $QH^-$ and proton from its C1-OH group shifted to E295. Note that shifting proton to E295 lowers the energy of the system but, in contrast to **R**, it is not followed by eT form $QH_2$ to $b_L^{ox}$. The relative energy (ΔE) for the structures is defined as kcal/mol differences from structure shown in A and was computed using def2-TZVP basis set in combination with PCM modeling protein environment defined by dielectric constant of 4 and radius probe of 1.4 Å. Data shown in blue frames represent spin populations gathered for the important fragments. Distances larger than 2.5 Å are not shown.

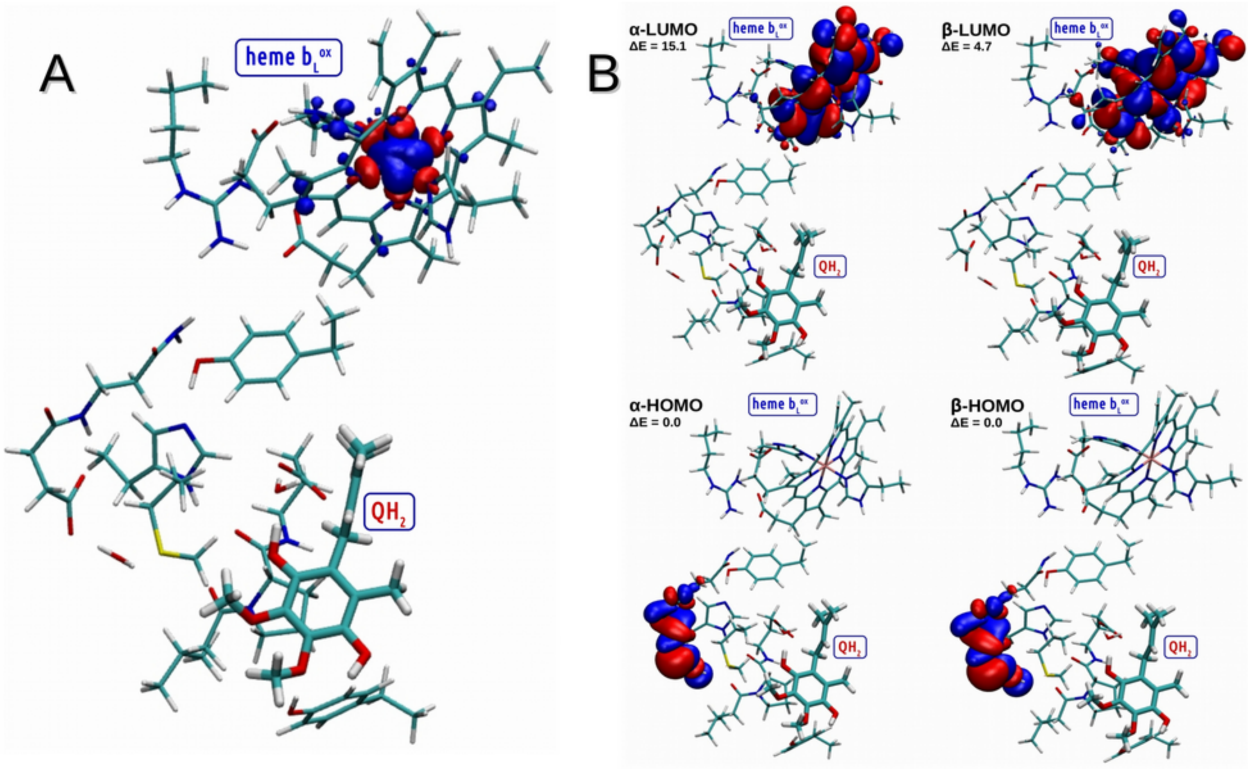


**Figure S4**: Distribution of spin frontier orbitals for structure shown in Fig. S3. A) Spin densities. B) Location of LUMO and HOMO.

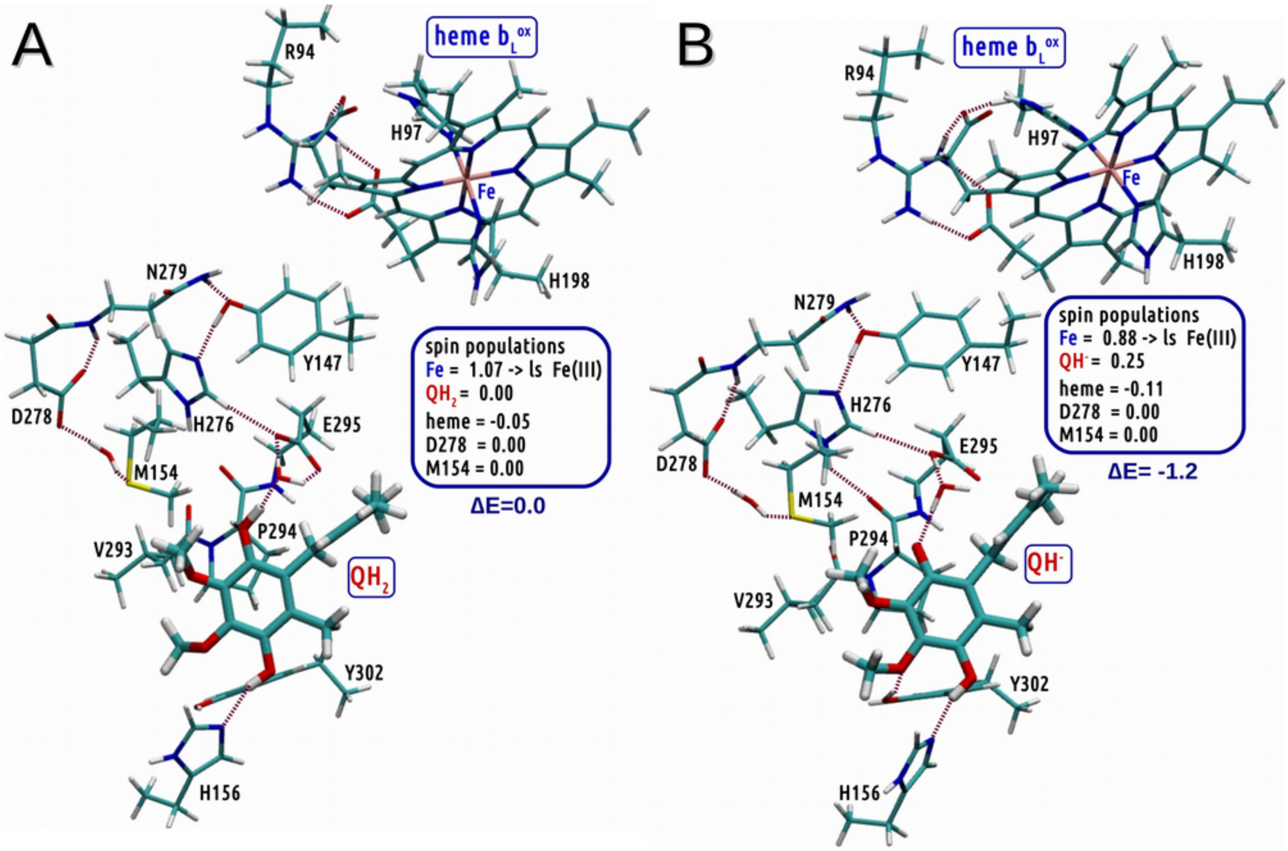


**Figure S5**: Optimized structures containing substrate, $b_L^{ox}$ and H156 ligand to 2Fe2S but without the cluster. A) The structure containing $QH_2$. B) The structure containing $QH^-$ and proton from its C1-OH group shifted to E295. Note that shifting proton to E295 lowers the energy of the system but, in contrast to **R**, it is not followed by eT form $QH_2$ to $b_L^{ox}$. The relative energy (ΔE) for the structures is defined as kcal/mol differences from structure shown in A and was computed using def2-TZVP basis set in combination with PCM modeling protein environment defined by dielectric constant of 4 and radius probe of 1.4 Å. Data shown in blue frames represent spin populations gathered for the important fragments. Distances larger than 2.5 Å are not shown.

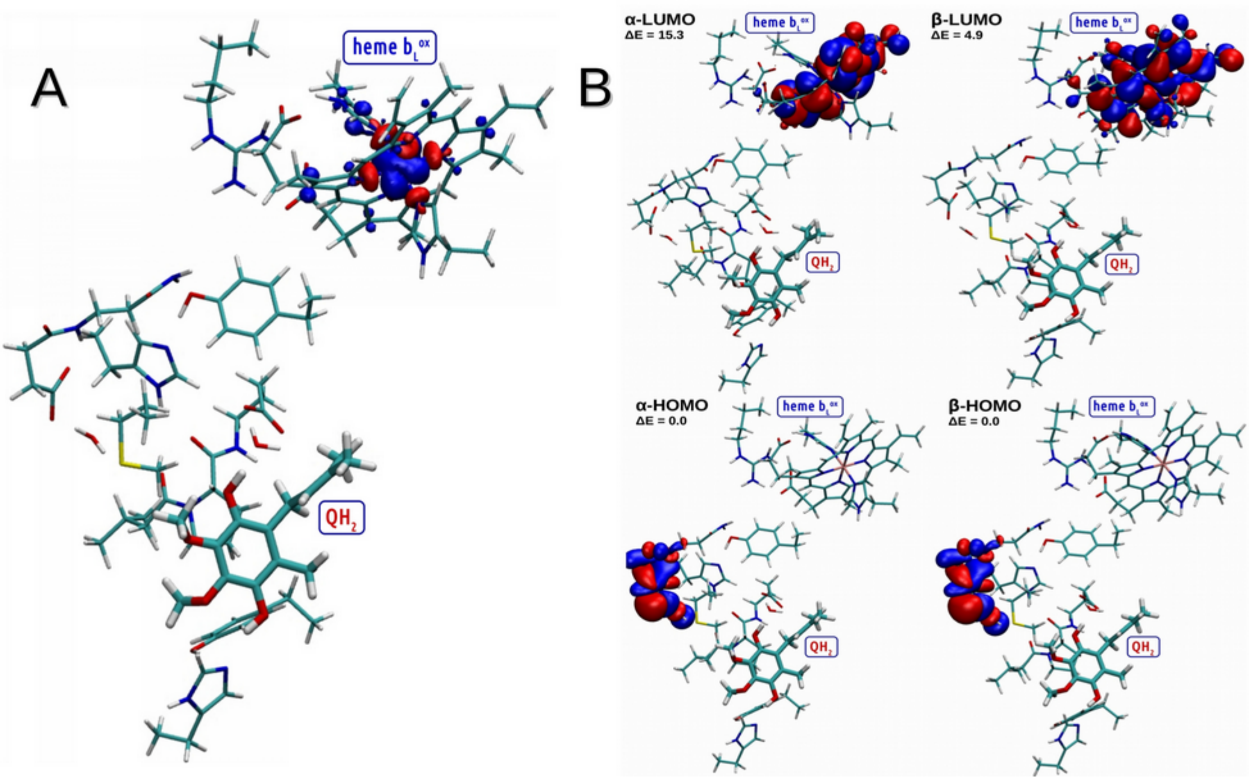


**Figure S6**: Distribution of spin frontier orbitals for structure shown in Fig. S5. A) Spin densities. B) Location of LUMO and HOMO.

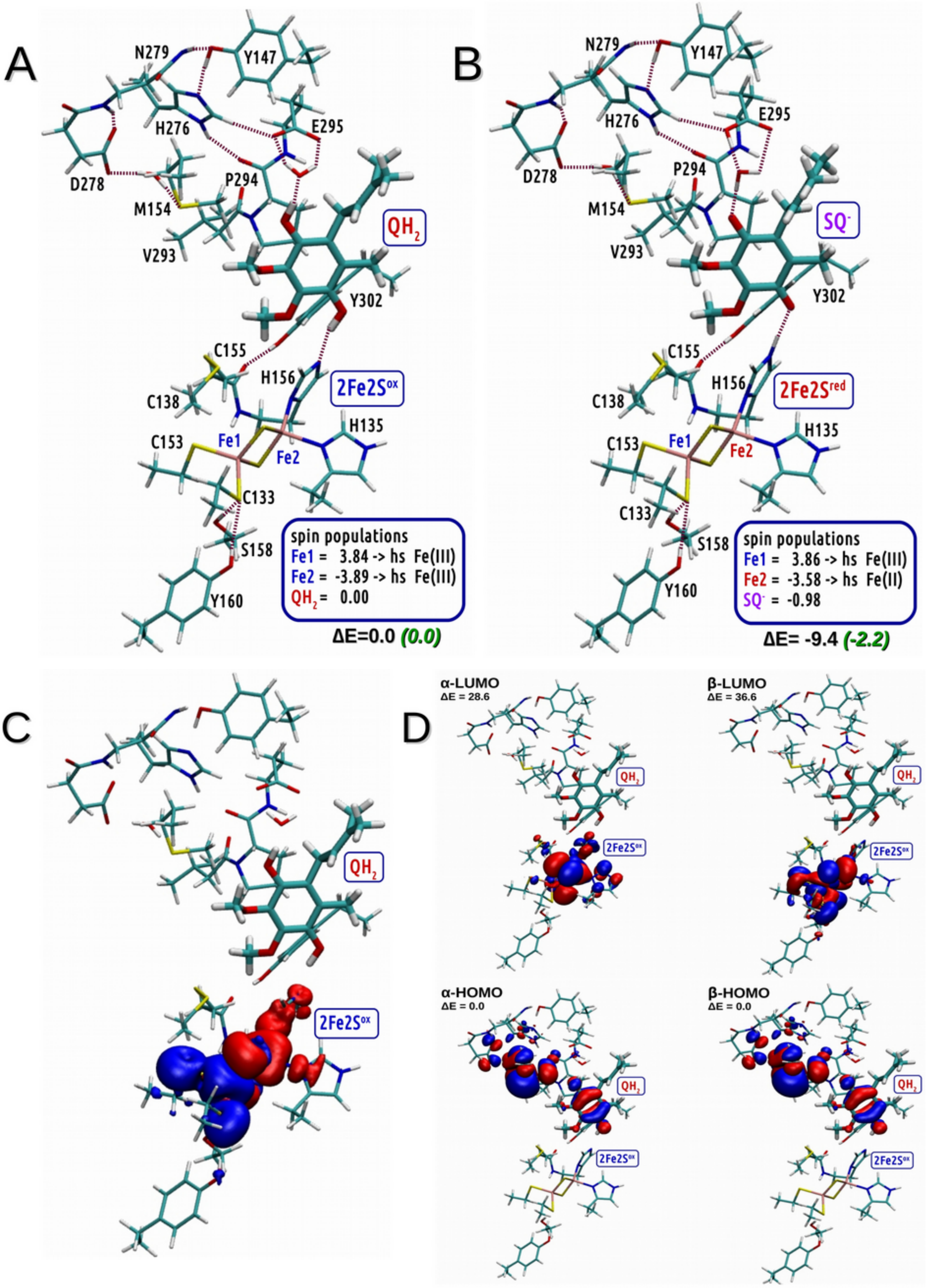


**Figure S7**: Structure **R** optimized without heme $b_L$, its ligands and residue R94. A) The structure containing $QH_2$. B) The structure containing $SQ^-$ and protonated E295 and H156. C) Spin densities for structure in A. D) Location of LUMO and HOMO orbitals for structure in A. The relative energy (ΔE) for the structures is defined as kcal/mol differences from structure shown in A and was computed using def2-TZVP basis set in combination with PCM modeling protein environment defined by dielectric constant of 4 and radius probe of 1.4 Å. Data shown in blue frames represent spin populations gathered for the important fragments. Distances larger than 2.5 Å are not shown.

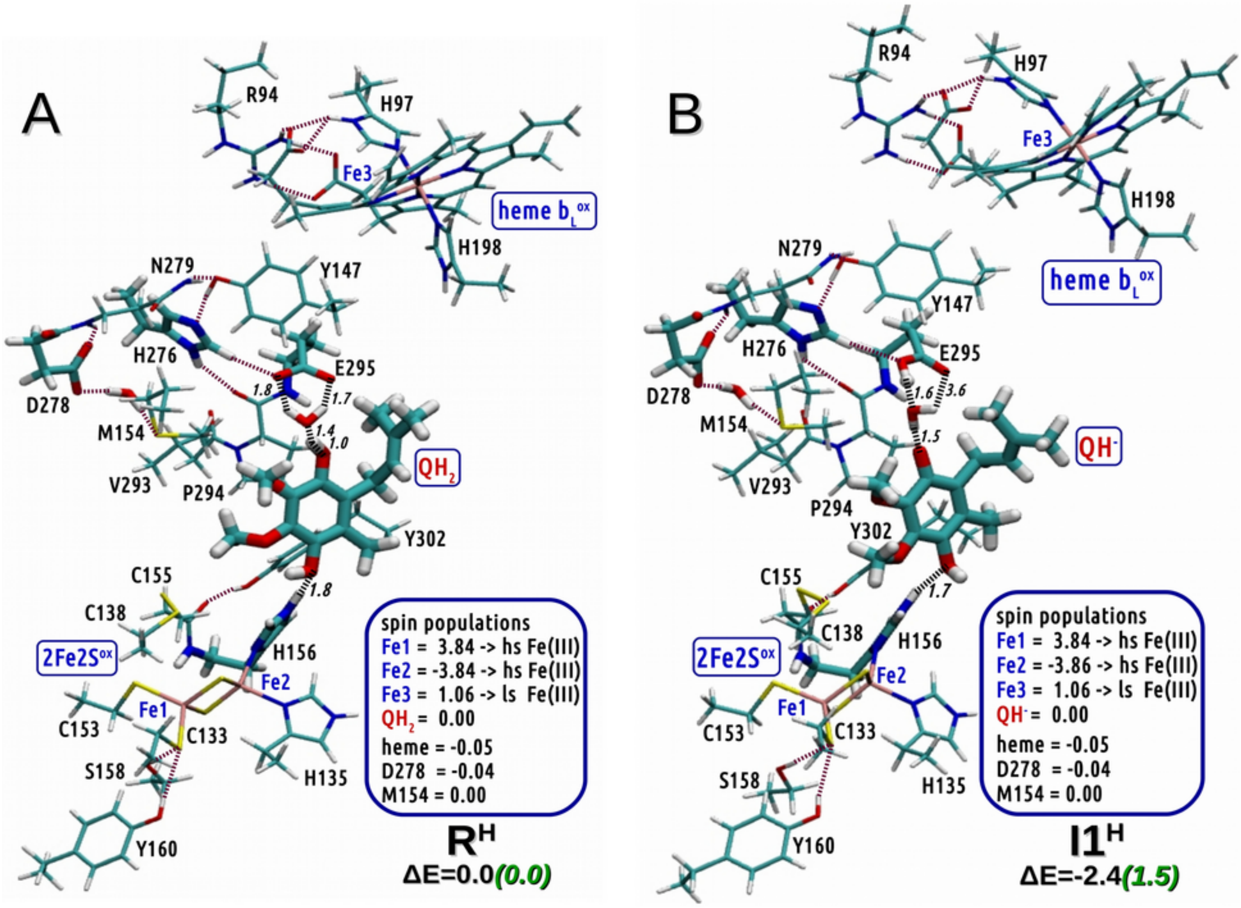


**Figure S8**: The structures containing $QH_2$ (**R$^H$**) and $QH^-$ (**I1$^H$**) complexes optimized with the model containing protonated H156. The QM results revealed that in this model the pT from $QH_2$ to E295 is not accompanied with eT. The relative energy (ΔE) for each of the structure is defined as kcal/mol differences from structure **R**. Black numbers in ΔE show relative energy computed with def2-SVP basis set in vacuum, while green numbers show respective energy using def2-TZVP basis set in combination with PCM modeling protein environment defined by dielectric constant of 4 and radius probe of 1.4 Å. Data shown in blue frames represent spin populations gathered for the important fragments. Distances larger than 2.5 Å are not shown.

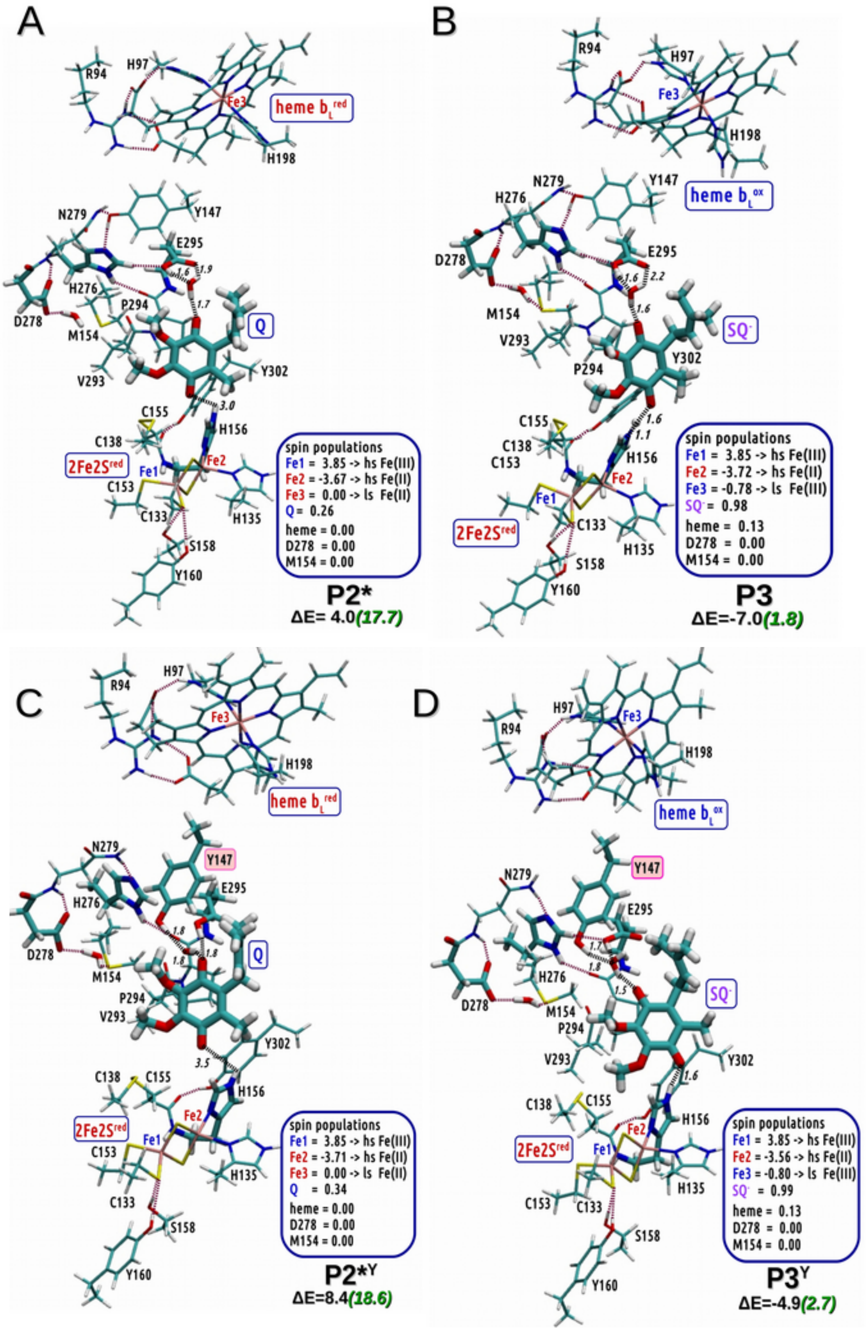


**Figure S9.** The optimized structures that evolve from **I2** under the conditions of blocked eT from $b_L$ to $b_H$. A) The structure **P2***, containing $b_L^{red}$, Q and 2Fe2S$^{red}$ resulting from elongation of the distance between H156 and SQ$^-$ from 1.5 (as in **I2**) to 3.0Å. Due to instability of this state, the geometry optimization required imposing a constraint on the distance between the C4 carbonyl oxygen of Q and the proton associated to $N_\tau$ nitrogen of H156. B) The structure **P3** containing $b_L^{ox}$ and SQ$^-$ ferromagnetically coupled to 2Fe2S$^{red}$. C) and D) shows the structures **P2***$^{Y}$ and **P3**$^{Y}$ that represent the counterparts of **P2*** and **P3**, respectively but with side chain of Y147 rotated toward Q. The relative energy (ΔE) for each of the structure is defined as kcal/mol differences from structure **R**. Black numbers in ΔE show relative energy computed with def2-SVP basis set in vacuum, while green numbers show respective energy using def2-TZVP basis set in combination with PCM modeling protein environment defined by dielectric constant of 4 and radius probe of 1.4 Å. Data shown in blue frames represent spin populations gathered for the important fragments. Distances larger than 2.5 Å are not shown.

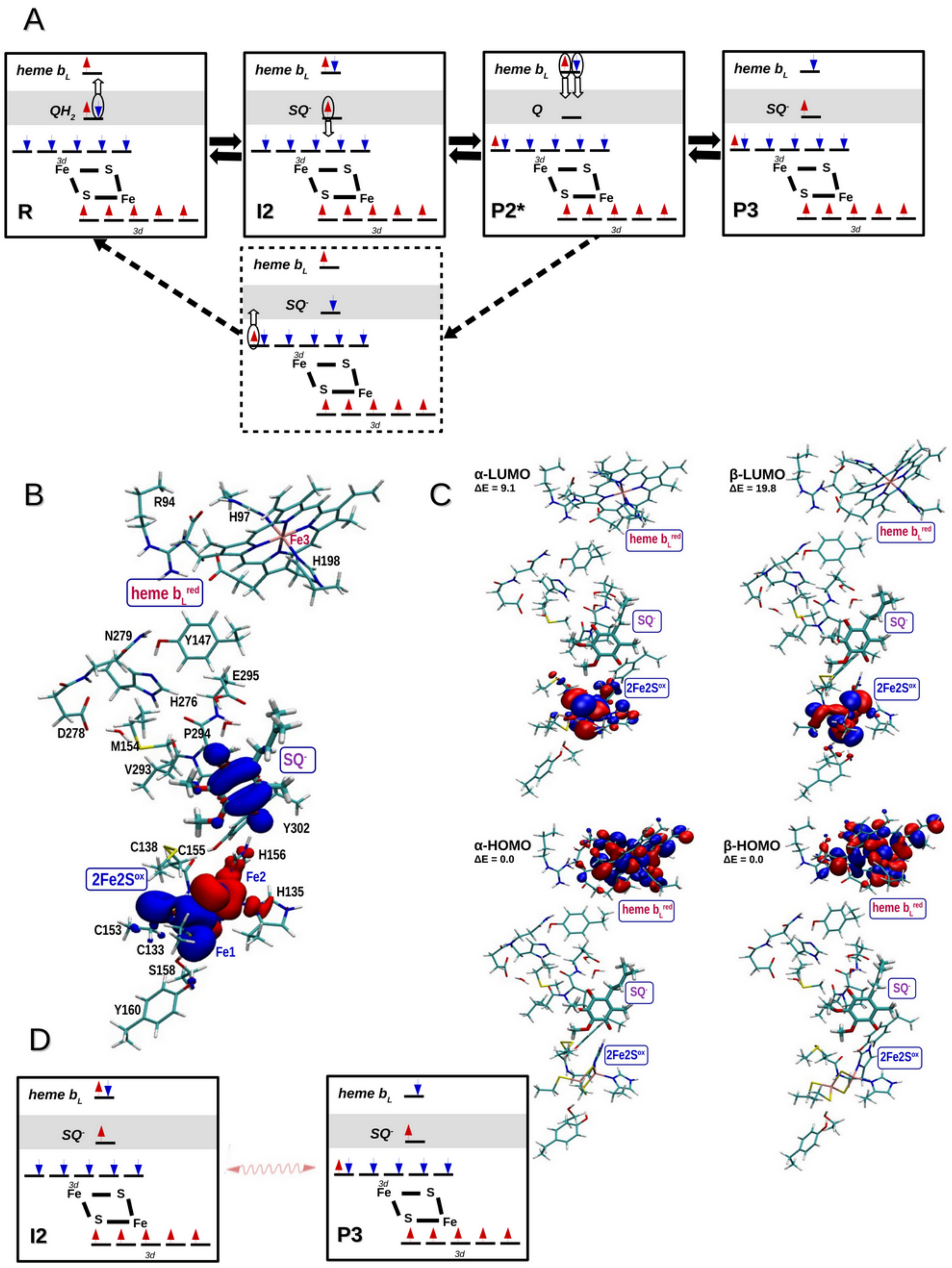


**Figure S10**. A) Schematic representation of the electron configurations of **R**, **I2**, **P2*** involved in the formation of **P3** state under conditions when eT from $b_L$ to $b_H$ is blocked. The states: **R** contains $b_L^{ox}$, $QH_2$ and 2Fe2S$^{ox}$; **I2** contains $b_L^{red}$, SQ$^-$ and 2Fe2S$^{ox}$; **P2*** contains $b_L^{red}$, Q and 2Fe2S$^{red}$; **P3** contains $b_L^{ox}$, SQ$^-$ ferromagnetically coupled to 2Fe2S$^{red}$. Red and blue arrows represent electrons with spin α and β, respectively. White arrows indicate a direction of eT leading to subsequent states. Solid and dashed frames show stable and unstable states, respectively. B) Spin density of optimized

**I2**. C) HOMO and LUMO orbitals for optimize **I2**. D) Scheme showing electron tunneling process (double wavy arrow) resulting from HOMO and LUMO orbitals of **I2** and **P3** configuration.

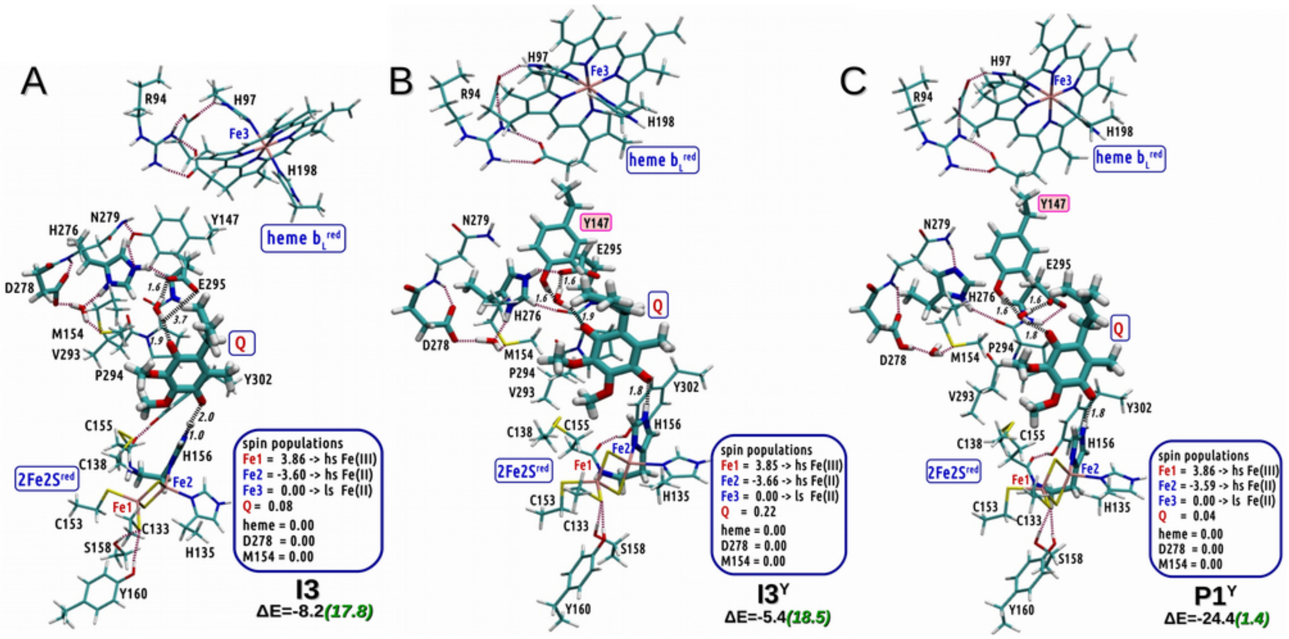


**Figure S11**. The most stable structures considered after formation of **I2** (pT from E295 to D278 via H276), under the conditions of blocked eT from $b_L$ to $b_H$. A) **I3** containing $b_L^{red}$-Q-2Fe2S$^{red}$ formed after pT from E295 to H276; B) **I3**$^{Y}$ containing $b_L^{red}$-Q-2Fe2S$^{red}$ formed after pT from E295 to H276 coupled with rotation of Y147 side chain; C) **P1**$^{Y}$ containing $b_L^{red}$-Q-2Fe2S$^{red}$ formed after pT from H276 to D278 and rotated side chain of Y147 interacting with quinone. The relative energy (ΔE) for each of the structure is defined as kcal/mol differences from structure **R**. Black numbers in ΔE show relative energy computed with def2-SVP basis set in vacuum, while green numbers show respective energy using def2-TZVP basis set in combination with PCM modeling protein environment defined by dielectric constant of 4 and radius probe of 1.4 Å. Data shown in blue frames represent spin populations gathered for the important fragments. Distances larger than 2.5 Å are not shown.

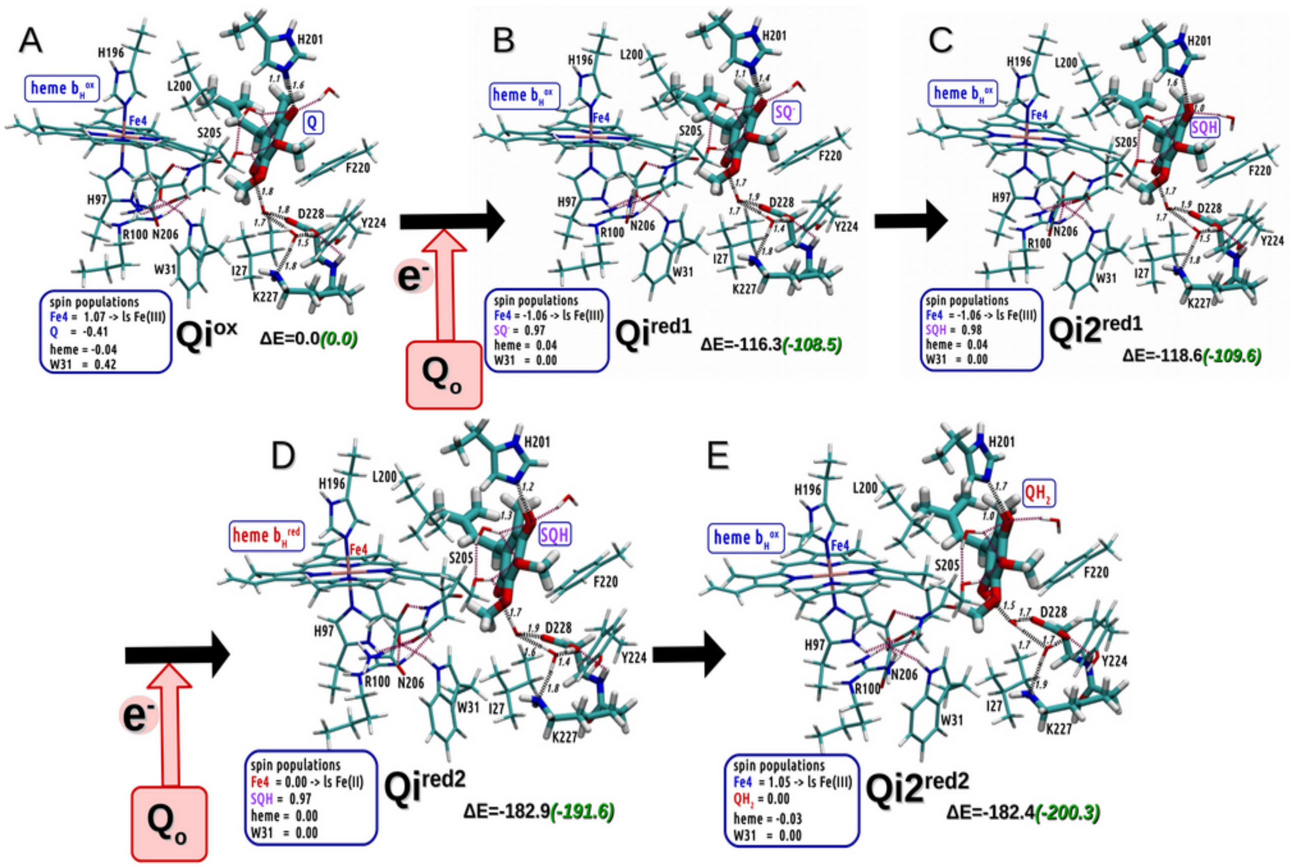


**Figure S12**. The optimized structures of the model encompassing $b_H$ and the $Q_i$ site. A) The state $Qi^{ox}$ contains oxidized $b_H$ and Q which is treated as reference for energy calculations. B) The structure $Qi^{red1}$ containing $b_H^{ox}$ and $SQ^{-}$. C) The structure $\mathbf{Qi2^{red1}}$ obtained from $\mathbf{Qi^{red1}}$ by pT from H201 to $SQ^{-}$. D) The structure $\mathbf{Qi^{red2}}$ which mimics the process of second eT from $b_L$ to $b_H$ under conditions in which the Qi site already contains SQ. E) The structure obtained from $\mathbf{Qi^{red2}}$ by pT from D228 to SQH which is coupled to eT from $b_H$ to SQH to create $QH_2$. The relative energy (ΔE) for each of the structure is defined as kcal/mol differences from structure $\mathbf{Qi^{ox}}$. Black numbers in ΔE show relative energy computed with def2-SVP basis set in vacuum, while green numbers show respective energy using def2-TZVP basis set in combination with PCM modeling protein environment defined by dielectric constant of 4 and radius probe of 1.4 Å. Data shown in blue frames represent spin populations gathered for the important fragments. Distances larger than 2.5 Å are not shown.

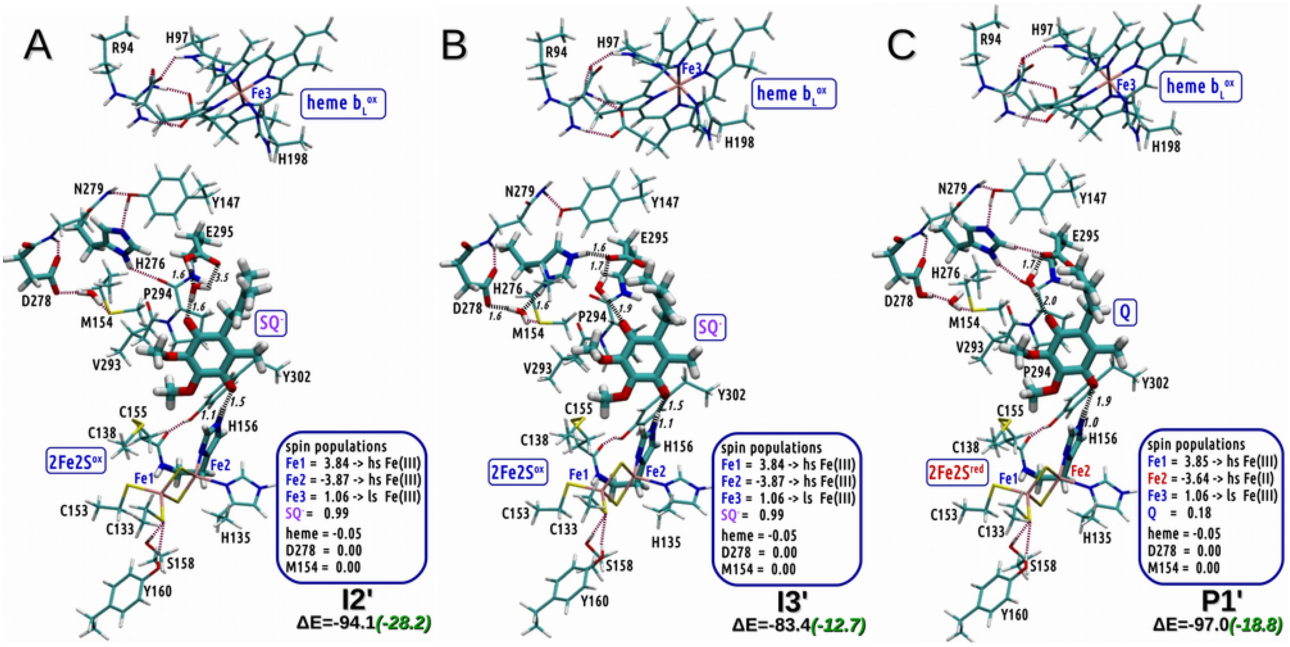


**Figure S13.** The most stable structures with Y147 rotated toward H276 obtained by removing the electron from $b_L$. A) The **I2'** structure containing $b_L^{ox}$, $SQ^-$ and 2Fe2S$^{ox}$. Energy of this state is respect to **I2** indicates how much energy is gained upon electron donation to $b_H$ and further to the $Q_i$ site. B) The **I3'** structure formed from **I2'** by pT from E295 to H276. C) The **P1'** structure formed from **I3'** by subsequent pT from H276 to D278. Note that this pT is coupled to eT from $SQ^-$ to 2Fe2S. As a result this state contains the product of EB: $b_L^{ox}$, Q, 2Fe2S$^{red}$. The relative energy (ΔE) for each of the structure is defined as kcal/mol differences from structure **R** corrected by the average energy released in the process of eT to the Qi site. Black numbers in ΔE show relative energy computed with def2-SVP basis set in vacuum, while green numbers show respective energy using def2-TZVP basis set in combination with PCM modeling protein environment defined by dielectric constant of 4 and radius probe of 1.4 Å. Data shown in blue frames represent spin populations gathered for the important fragments. Distances larger than 2.5 Å are not shown.

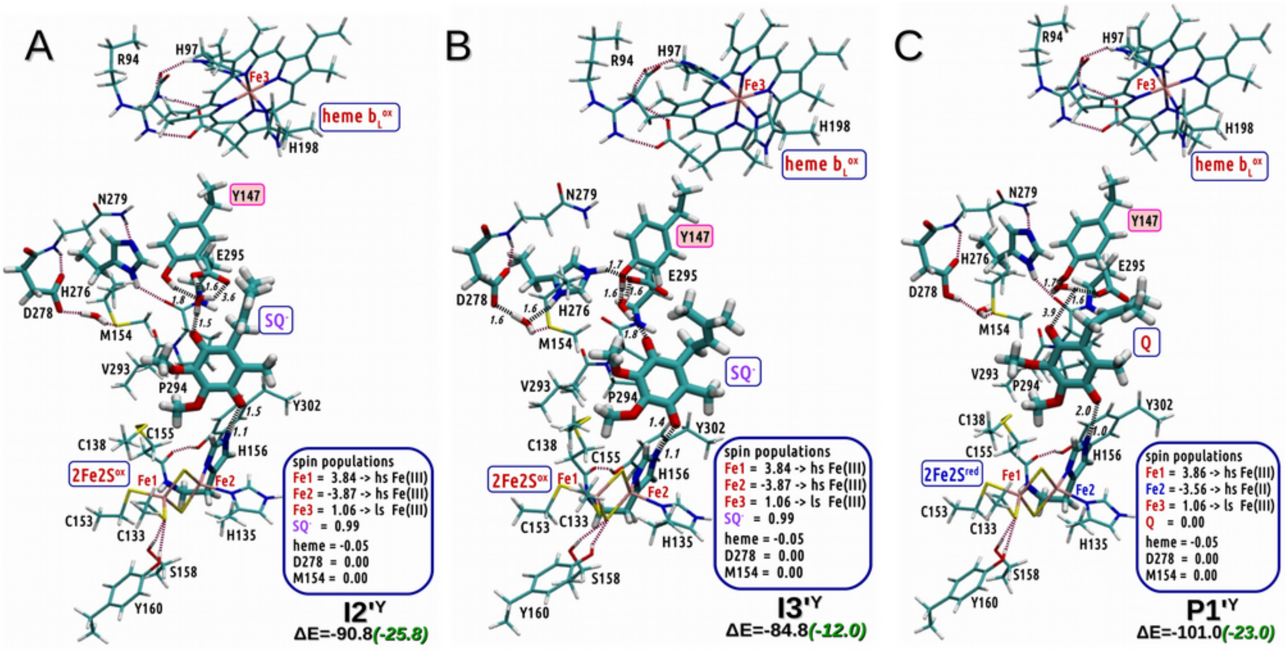


**Figure S14**. Counterpart structures to the structures shown in figure S15 but with Y147 rotated toward the ring of Q or SQ. A) The **I2'**$^{Y}$ counterpart of **I2'**. B) The **I3'**$^{Y}$ counterpart of **I3'**. C) The **P1'**$^{Y}$ counterpart of **P1'**. The most stable structures with Y147 rotated toward H276 obtained by removing the electron from $b_L$. The relative energy (ΔE) for each of the structure is defined as kcal/mol differences from structure **R** corrected by the average energy released in the process of eT to the Qi site. Black numbers in ΔE show relative energy computed with def2-SVP basis set in vacuum, while green numbers show respective energy using def2-TZVP basis set in combination with PCM modeling protein environment defined by dielectric constant of 4 and radius probe of 1.4 Å. Data shown in blue frames represent spin populations gathered for the important fragments. Distances larger than 2.5 Å are not shown.

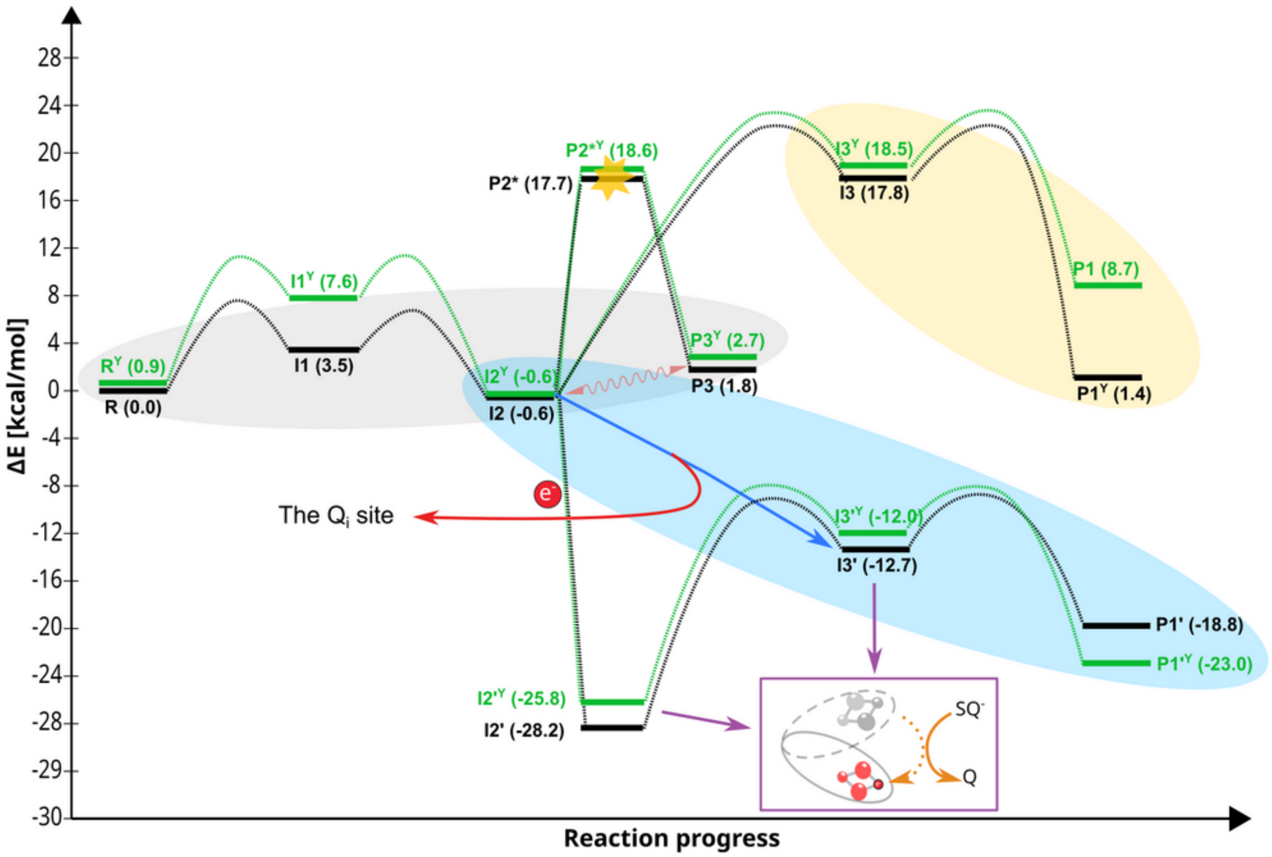


**Figure S15. Energy levels of the optimized states considered in the DFT calculations of the EB process.** Solid horizontal black and green lines represent the energy levels of states corresponding to two conformations of the Y147 residue: shifted outward or oriented toward the quinone ring, respectively. Dotted lines depict transitions between these states. The blue arrow indicates the average downhill reaction associated with electron transfer eT from $b_L$ to the $Q_i$ site (red arrow), coupled with proton transfer. The yellow star in **P2*** and **P2*ʸ** marks an unstable state whose energy was estimated using optimization involving geometric constrains. The magenta arrow highlights the motion of the ISP-HD domain away from the $Q_o$ site (magenta box), leading to eT from $SQ^-$ to the 2Fe2S cluster. This motion may be further driven by energy release from eT toward the Qi site and induces eT from $SQ^-$ to 2Fe2S. If eT from $b_L$ to $b_H$ is inhibited (e.g., in the presence of antimycin), the reaction becomes trapped in the states enclosed by the gray oval. Furthermore, reaching the final product states (**P1ʸ** and **P1**) would require traversing the pathway indicated by the yellow oval, which is highly improbable due to large energy barriers and the lack of energetic gain. Conversely, when eT from $b_L$ to the $Q_i$ site is unimpeded, the reaction proceeds to the product states (**P1'** and **P1'ʸ**), involving the states marked by the blue oval. It is noteworthy that eT from $b_L$ to $b_H$ is strongly coupled with proton transfer (pT) from E295 to H276. Therefore, the states **I2'ʸ** and **I2'** are likely transient and may not form during uninhibited EB reactions.
